# Epithelial cells of the intestine acquire cell-intrinsic inflammation signatures during ageing

**DOI:** 10.1101/2021.12.19.473357

**Authors:** Maja C. Funk, Jan G. Gleixner, Florian Heigwer, Erica Valentini, Zeynep Aydin, Elena Tonin, Jenny Hetzer, Danijela Heide, Oliver Stegle, Mathias Heikenwalder, Michael Boutros

## Abstract

During ageing, cell-intrinsic and extrinsic factors lead to the decline of tissue function and organismal health. Disentangling these factors is important for developing effective strategies to prolong organismal healthspan. Here, we addressed this question in the mouse intestinal epithelium, which forms a dynamic interface with its microenvironment and receives extrinsic signals affecting its homeostasis and tissue ageing. We systematically compared transcriptional profiles of young and aged epithelial cells *in vivo* and *ex vivo* in cultured intestinal organoids. We found that all cell types of the aged epithelium exhibit an inflammation phenotype, which is marked by MHC class II upregulation and most pronounced in enterocytes. This was accompanied by elevated levels of the immune tolerance markers PD-1 and PD-L1 in the aged tissue microenvironment, indicating dysregulation of immunological homeostasis. Intestinal organoids from aged mice still showed an inflammation signature after weeks in culture, which was concurrent with increased chromatin accessibility of inflammation-associated loci. Our results reveal a cell-intrinsic, persistent inflammation phenotype in aged epithelial cells, which might contribute to systemic inflammation observed during ageing.

## Introduction

During ageing, cell-intrinsic and extrinsic factors act in concert and lead to the functional decline of tissues and organs ^1,2^. An organ that is particularly exposed to external signals is the intestine. In homeostasis, the intestine forms a dynamic interface with the microenvironment and serves as a protective barrier against external stimuli ^3–5^. Furthermore, the intestinal epithelium balances immune response between effective clearance of invading pathogens and tolerance against the commensal microbiome ^4^and ensures sufficient nutrient uptake over the entire lifetime of an organism.

To maintain tissue homeostasis the small intestine relies on highly proliferative stem cells to continuously renew the entire epithelium. Intestinal stem cells (ISCs) reside in their stem cell niche, called crypt. New progenitor cells leave the crypt, enter the transit-amplifying zone, and are pushed upwards to the villi, the functional units of the intestine ^6^. Along the crypt-villus axis, different cell types can be found: ISCs, absorptive enterocytes, secretory cells, including Paneth cells, Goblet cells, Enteroendocrine cells, and chemosensory Tuft cells ^7^. Paneth cells, intercalated between the ISCs, are part of the stem cell niche where they secrete growth factors essential for the stem cells and antimicrobial peptides as a first-line defense against microbes ^8^. Additionally, immune cells and the microbiome in the intestinal microenvironment can directly regulate ISC function and epithelial homeostasis ^9–11^.

Upon ageing, intestinal stem cell capacity is impaired ^12–14^, in part due to reduced canonical Wnt signaling in ISCs, neighboring Paneth cells, and the subepithelial mesenchyme ^12,14^. Additionally, the intestinal epithelium is exposed to extrinsic factors from the microenvironment, which are changing with age ^5^. During ageing, direct interaction with immune cells and the microbiome causes inflammatory responses in the aged intestine ^15–17^. However, it remains unclear which ageing phenotypes establish over time within epithelial cells and persist independently from external signals from the microenvironment.

To identify molecular features of ageing in the intestinal epithelium, we profiled transcriptional changes of the aged intestinal epithelium in bulk and in single cells, to map ageing signatures to individual cell types of the intestine *in vivo*. We show that the age-related chronic inflammation phenotype, termed inflammaging ^18^, is marked in the intestine by upregulation of MHC class II molecules in all epithelial cell types and is most pronounced in enterocytes. Using *ex vivo* intestinal organoid cultures, we identified epithelium-intrinsic inflammaging signatures that persist in culture, independent of external signals from the microenvironment. Analysis of open chromatin accessibility in organoids further revealed age-related chromatin remodeling in inflammation-associated loci. In the aged intestinal microenvironment, we detected elevated levels of the immune tolerance markers PD-1 and PD-L, indicating T cell exhaustion and dysregulation of immunological homeostasis upon ageing. In summary, we reveal cell-intrinsic inflammaging signatures in the aged intestinal epithelium, which provides evidence that epithelial cells might contribute to systemic inflammation and disbalance of immune homeostasis during ageing.

## Results

### Inflammation of the ageing intestine is marked by MHC class II upregulation

In this study, we set out to dissect ageing signatures of the intestinal epithelium that are dependent or independent of direct external signals, such as from immune cells or the microbiome. To this end, we profiled transcriptional changes in the aged intestinal epithelium from freshly isolated epithelial cells and compared them to transcriptional changes in *ex vivo* intestinal organoids from young and aged mice. Organoid cultures are devoid of other non-epithelial cells and allow to identify intrinsic ageing phenotypes that persist independently of direct signals from the surrounding microenvironment.

To first investigate the *in vivo* ageing effect in all the intestinal cell types, we performed bulk and single-cell RNA-sequencing (RNA-Seq) from freshly isolated epithelial cells (EpCam^+^, CD45^-^) of the proximal small intestine of three young and aged mice each (Fig 1a). In a principal component analysis (PCA), we found that the age of the mice was the first driving force to discriminate the samples of the bulk RNA-Seq experiment (Extended Data Fig 1a). We identified 508 genes significantly differentially expressed upon ageing, of which 190 genes were upregulated and 318 genes were downregulated (Fig 1b, Extended Data Fig 1b). Downregulated genes included alpha-defensins (*Defa3, Defa17, Defa24, Defa26*) (Fig 1b, Extended Data Fig 1c), which are specifically secreted by Paneth cells for anti-microbial defense ^8^. This downregulation indicates reduced functionality of Paneth cells and mucosal immunity upon ageing. We also detected downregulation of Wnt3 in the aged epithelium, consistent with previous reports (Fig 1b) ^12,14^. Upregulated genes comprised various immune response and interferon-inducible genes, such as *Igtp, Ly6e, Ccr2, Ifitm3, P2rx7, and Ifi47* (Fig 1b, 1c). We also found increased expression for genes encoding for interferon-inducible guanylate-binding proteins (Gbps) (*Gbp2, 6, 7, 8, 9*), which are involved in inflammasome activation ^19^ (Extended Data Fig 1d). Most notably, we identified several genes involved in Class II major histocompatibility complex (MHC class II)-mediated antigen presentation highly upregulated in the aged intestinal epithelium (Fig 1d), which included the invariant chain *Cd74* and the MHC class II transactivator *Ciita* (Fig 1c).

**Fig 1:**
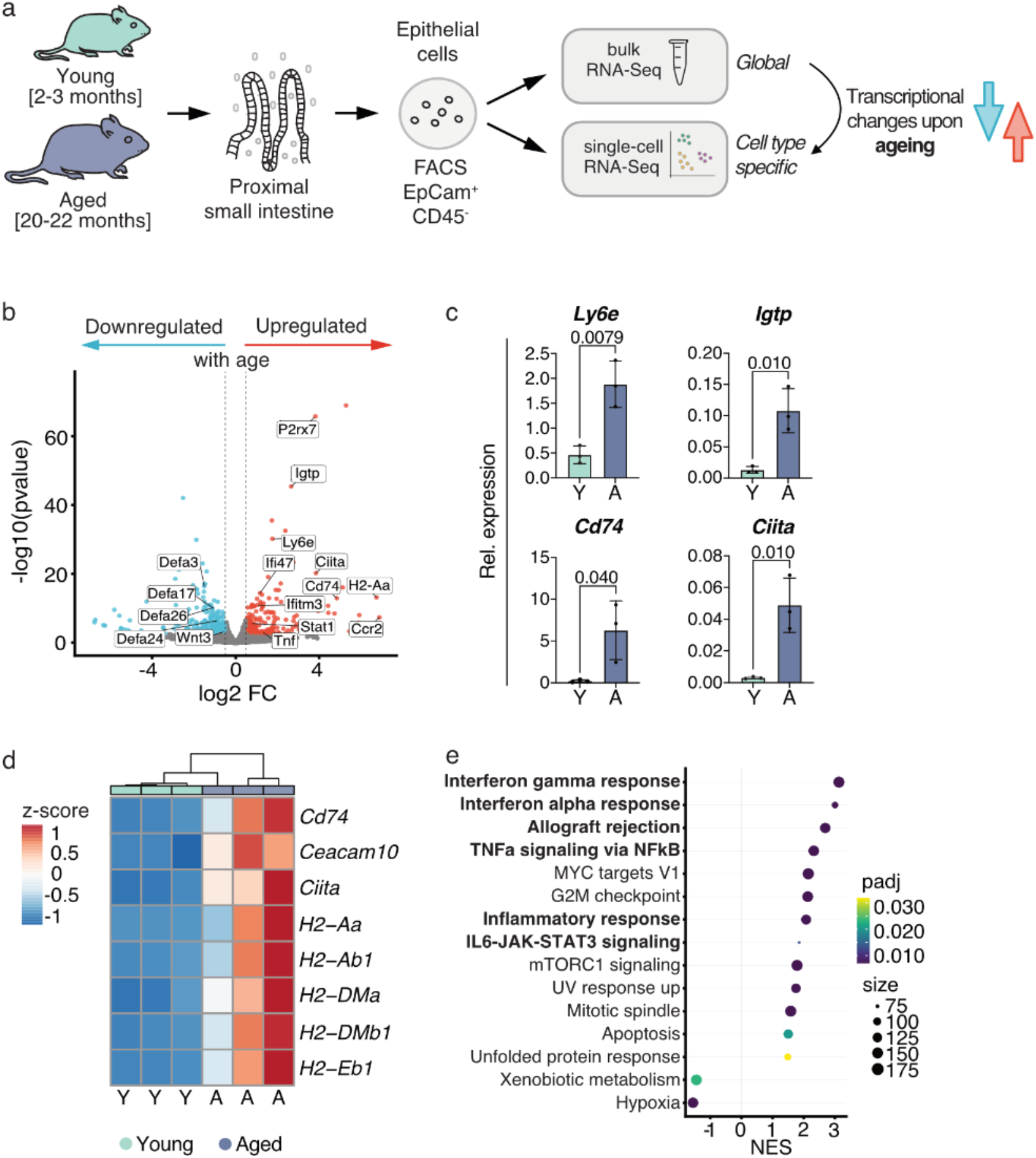
Inflammation of the ageing intestine is marked by MHC class II upregulation. **a)** Schematic of the experimental approach of bulk and single-cell RNA-Seq of *in vivo* epithelial cells from the proximal small intestine of young (2-3 months of age) and aged (20-22 months of age) mice. **b)** Immune-related genes are upregulated in the aged intestinal epithelium. Volcano plot showing differentially expressed genes upon ageing (aged over young) in the small intestinal epithelium, red: significantly (FDR≤10%) upregulated (log2 FC > 0.5), blue: significantly (FDR≤10%) downregulated (log2 FC < (-0.5)), grey: not significant and absolute log2 FC < 0.5. **c)** *Ly6e, Igtp, Cd74*, and *Ciita* are upregulated in the aged small intestinal epithelium. Transcript levels were assessed in young and aged small intestinal epithelium by qRT-PCR (normalized to *36B4*) (n=3 mice per age); statistical significance was tested by an unpaired t-test (two-sided) and p-Value is indicated on respective comparison; error bars indicate mean with standard deviation. **d)** Upregulation of MHC II gene expression upon ageing in the small intestinal epithelium. Relative mean expression as z-score per MHC II gene (row) and sample (column) (3 young, indicated by “Y”, 3 aged, indicated by “A”), color code indicates z-score. **e)** Aged intestinal epithelium is enriched for Interferon signaling and inflammatory response signatures. Normalized enrichment score (NES) based on gene set enrichment analysis on hallmark gene sets (MSigDB), filtered for (FDR≤5%), dot size indicates the size of the respective gene set, color code indicates FDR (log2 FC: log_2_ fold change).

In order to gain functional insights on the ageing phenotype of the intestine, we performed gene set enrichment analysis (GSEA) on the bulk RNA-Seq data. This analysis revealed enrichment for Interferon-gamma (IFN*γ*) and -alpha (IFNα) response signatures as well as other inflammation-associated signaling pathways, including allograft rejection, TNFα signaling via NF*κ*B, inflammatory response and IL6-JAK-STAT3 signaling (Fig 1e). Low chronic inflammation upon ageing is defined as inflammaging and we collectively defined these six inflammation-associated gene sets as inflammaging gene set. Beyond the inflammaging gene set, we detected enrichment for additional ageing-associated features, including mTORC1 signaling, UV response, and unfolded protein response. Moreover, xenobiotic metabolism, involved in detoxification, was downregulated with age (Fig 1e).

In summary, our bulk RNA-Seq approach demonstrated that the aged intestinal epithelium shows signs of reduced mucosal immunity and a pronounced inflammaging signature, which is marked by the upregulation of MHC class II genes. Of note, most of the inflammation and immune-related genes upregulated during ageing already showed a basal, low-level expression at a young age. This fact highlights the intestinal epithelium as an immune-responsive tissue and suggests that immune-responsiveness becomes unbalanced during ageing.

### Intestinal inflammaging is driven by transcriptional changes within individual cell types

Next, we aimed to assess if the inflammaging signature observed in bulk is primarily driven by transcriptional changes within individual cells and cell types or by cell type ratio shifts. To this end, we isolated and sequenced single epithelial cells (EpCam^+^, CD45^-^) from the small intestine of three young and aged mice (Fig 1a). Transcriptomes with more than 10 000 total counts and less than 15 % of counts from mitochondrial genes passed our quality control, resulting in 13 360 high-quality transcriptomes (8 240 from young and 5 140 from aged mice) (Extended Data Fig 2). To assign a cell type to each of our single-cell transcriptomes, we transferred refined cell type labels from a reference data set ^20^ independently for each replicate. This approach distinguished all known intestinal cell types (Fig 2a, Extended Data Fig 3a-f) at similar fractions as reported before ^20^ (Extended Data Fig 4a). Importantly, all identified cell type clusters corresponded to epithelial cell types, strongly indicating that our results were not confounded by non-epithelial cells, such as immune cells. PCA for all cell types together showed that cell type identity was largely maintained upon ageing (Extended Data Fig 4b). While within individual cell types the age of the mice displayed as the major source of variance between the samples (Extended Data Fig 4c-f), demonstrating robust transcriptomic changes in the aged epithelium down to a single cell type level. In a tSNE embedding, young and aged cells were distributed evenly across the different clusters (Fig 2b, Extended Data Fig 4g), and the quantification of the fraction of cells per cell type did not reveal significant differences upon ageing (Fig 2c).

**Fig 2:**
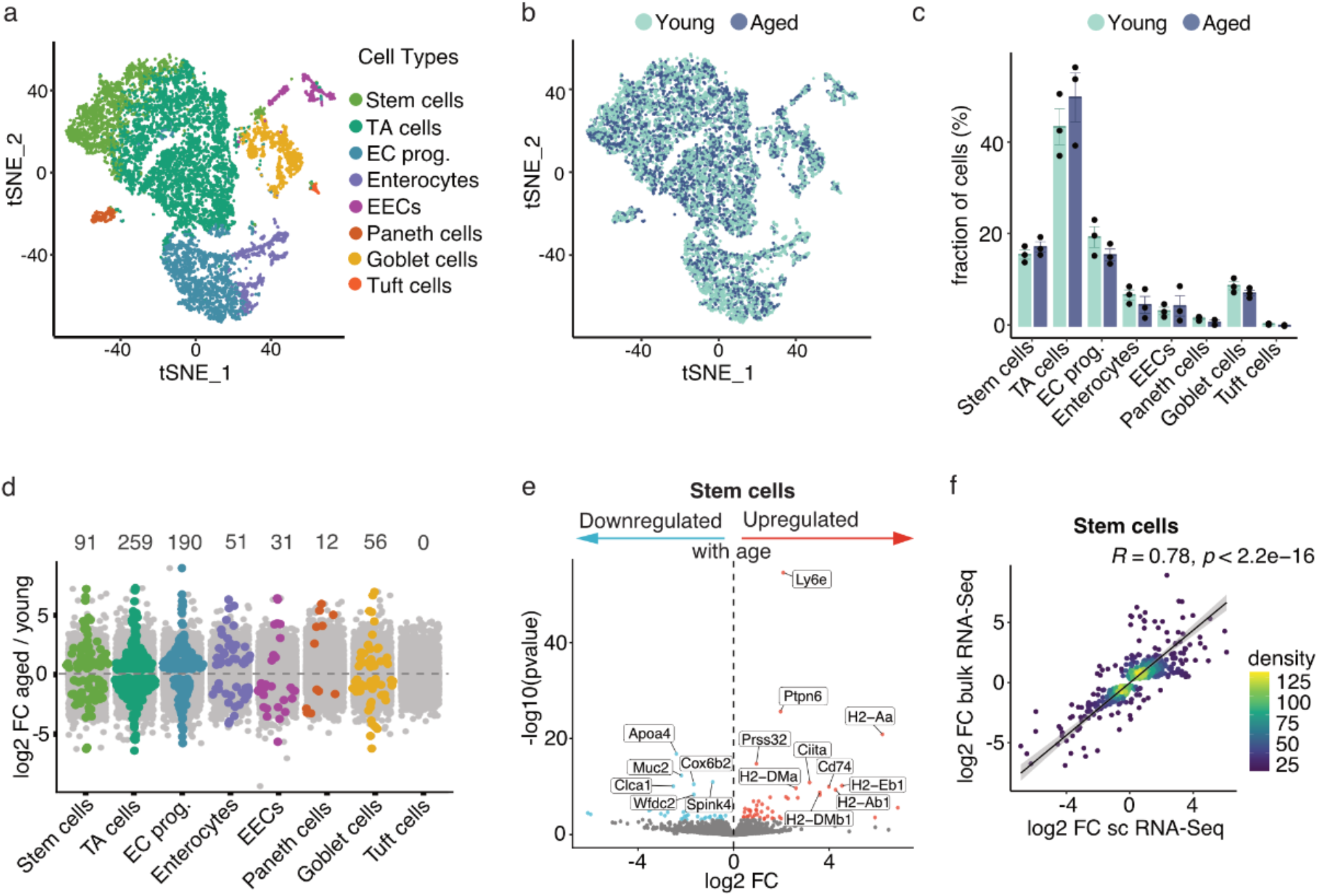
Intestinal inflammaging is driven by transcriptional changes within individual cell types. **a)** Cell type assignment of all 13 360 sequenced cells from six different mice (3 young, 3 aged) visualized in a tSNE embedding. Cells are colored based on their cell type assignment **b)** tSNE visualization of all 13 360 sequenced cells, cells are color-coded by the age of the mice they originated from. **c)** Cell type ratios of the small intestine are stable upon ageing. Average fraction of cells (in %) per sample for all young and all aged samples; data points indicate the percentage of individual samples. **d)** Number of differentially expressed genes in the small intestinal epithelium between young and aged per cell type. Genes upregulated upon ageing are displayed as log2 FC > 0 (aged/young), genes downregulated upon ageing are displayed as log2 FC < 0. Significantly (FDR≤10%) differentially expressed genes are colored in the respective cell type-specific color, not significantly changed genes are displayed in grey. **e)** Volcano plot showing differentially expressed genes upon ageing (aged/young) in the intestinal stem cells, red: significantly (FDR≤10%) upregulated (log2 FC > 0), blue: significantly (FDR≤10%) downregulated (log2 FC < (−0.5)), grey: not significant. The 20 most significantly differentially expressed genes are labeled and include several MHC class II encoding genes. **f)** Significantly differentially expressed genes in the aged intestinal epithelium identified by bulk RNA-Seq (y-axis) correlate with those detected in intestinal stem cells by single-cell RNA-Seq (x-axis). Genes are colored by the density of points in the respective plot area. A Deming regression line, as well as the Pearson correlation coefficient R and a p-value (p) of a test for correlation, are shown in black (log2 FC: log_2_ fold change, TA: transit-amplifying, EC prog.: enterocyte progenitors, EECs: enteroendocrine cells).

To compare the transcriptional changes upon ageing in individual cell types with our bulk analysis, we assessed differentially expressed genes (DEGs) for each cell type (Fig 2d). We identified significant DEGs in all cell types, except in Tuft cells, due to the low number of sequenced cells leading to insufficient statistical power. With 259 genes, we detected the highest number of DEGs in transit-amplifying (TA) cells. In intestinal stem cells (ISCs), 91 genes were significantly changed upon ageing. Among these DEGs, we identified the interferon-inducible *Ly6e* as the most significantly upregulated gene in aged ISCs and many MHC class II genes, including *Cd74* and *Ciita* (Fig 2e). These inflammation-associated genes overlapped with results obtained from bulk RNA-Seq (Fig 1c,d), and further analysis revealed a strong correlation between DEGs identified in bulk tissue and in the stem cell cluster (Fig 2f). Of note, we found that transcriptional changes upon ageing in all the individual cell types of the intestinal epithelium correlated with the DEGs observed in bulk to varying degrees (Extended Data Fig 5a-h).

Thus, profiling transcriptional changes within individual cell types of the aged intestinal epithelium unveiled robust changes upon ageing on a single cell and cell type level, which included inflammation-associated genes, such as *Ly6e* and MHC class II. Correlation analysis further corroborated our hypothesis that the inflammaging signature is primarily driven by transcriptional changes within individual cell types of the aged intestinal epithelium.

### Inflammaging signature is established in all cell types of the aged intestinal epithelium

To determine which cell types contribute to the inflammaging signature of the aged intestine and to gain functional insights on the ageing phenotype of every cell type, we performed GSEA based on the DEGs in each individual cell type cluster (Fig 3a).

**Fig 3:**
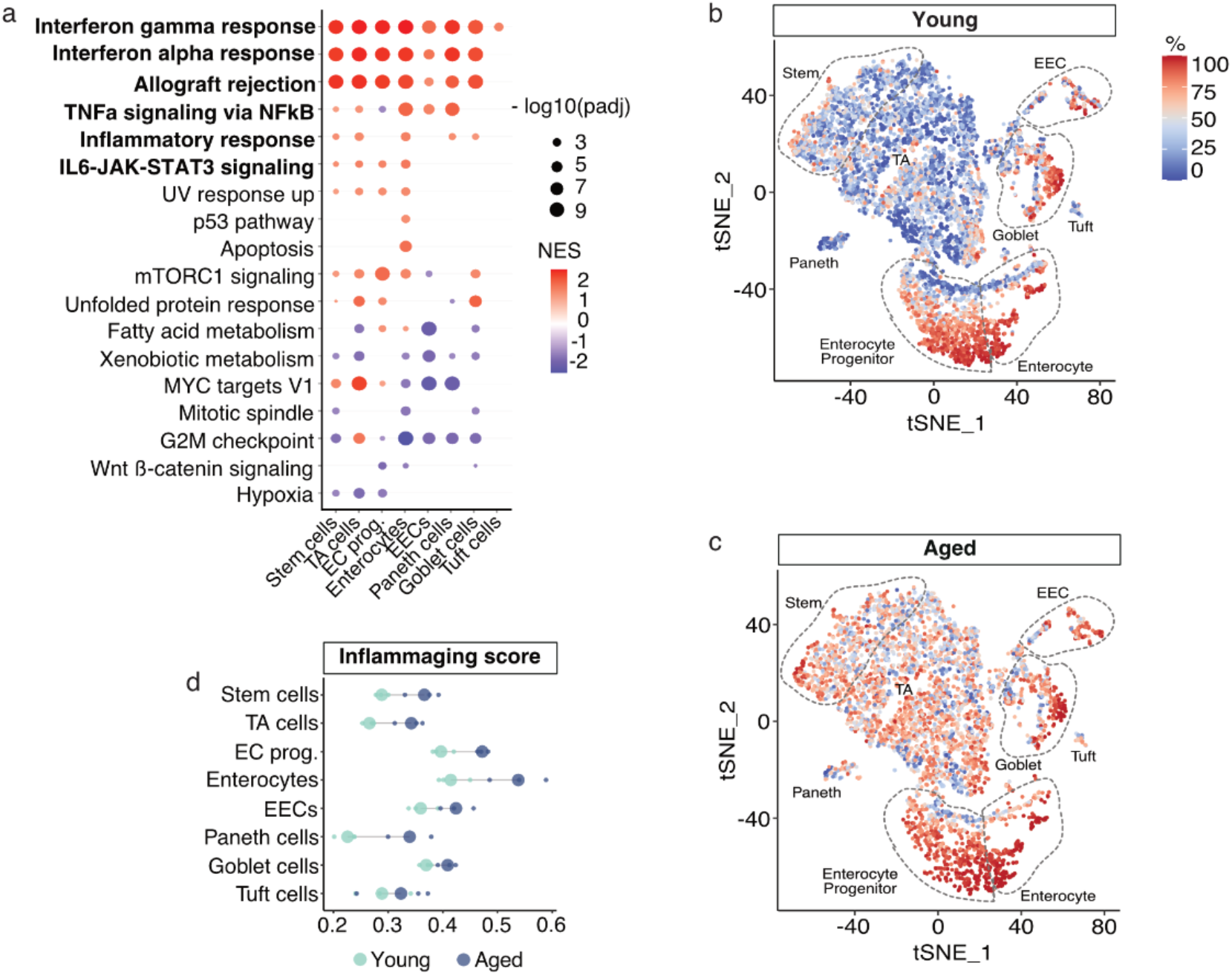
Inflammaging signature is established in all cell types of the aged intestinal epithelium. **a)** Gene set enrichment analysis reveals broad enrichment of inflammaging gene sets in all cell types. Normalized enrichment score (NES) based on gene set enrichment analysis on hallmark gene sets (MSigDB) for all cell types, filtered for (FDR≤5%), dot size indicates significance value (FDR), color code indicates NES. A list of gene sets was selected according to the list in Fig 1d. **b** & **c)** tSNE visualization of all **b)** young and **c)** aged cells separate and color-coded for their individual inflammaging score (quantile normalized). **d)** Inflammaging score calculated for each cell type and for each sample (3 young and 3 aged mice) demonstrates an increased inflammaging score for all cell types and most pronounced in enterocytes (TA: transit-amplifying, EC prog.: enterocyte progenitors, EECs: enteroendocrine cells).

This analysis revealed an IFN*γ* response signature in all cell types, including Tuft cells, while IFNα response signature and allograft rejection were enriched in all but Tuft cells. The other gene sets of the previously defined inflammaging signature (TNFα signaling via NF*κ*B, inflammatory response, IL6-JAK-STAT3 signaling) were also enriched across multiple cell types. At the same time, enterocytes were the only cell type that showed enrichment for all six gene sets. To quantify the inflammaging signature for every cell type, we calculated an *inflammaging score* per cell. For this, we aggregated the expression levels for all genes of the inflammaging gene set and compared them to a random set of 100 genes with similar expression levels. We plotted the score to every single cell for the young (Fig 3b) and aged (Fig 3c) intestine separately in a tSNE visualization, which clearly illustrated an increased inflammaging signature in all cell types of the intestinal epithelium upon ageing. Aggregation of the *inflammaging score* for each sample and each cell type corroborated that it was increased with age in all cell types (Fig 3d), while enterocytes showed the highest score overall, as indicated by GSEA (Fig 3a).

In summary, our analysis demonstrated that the inflammaging signature is present in all cell types of the aged intestinal epithelium and is most pronounced in enterocytes.

### *Ex vivo* intestinal organoids reveal an epithelium-intrinsic inflammaging signature

Inflammatory responses in the intestine, for example during inflammatory bowel disease (IBD), often arise from crosstalk between the epithelial cells, the immune system, and the gut microbiome ^4^. To investigate whether the inflammaging signature of the aged intestinal epithelium relies entirely on direct interaction with surrounding cells of the microenvironment, we took advantage of the 3-dimensional organoid culture system. Small intestinal organoids are *ex vivo* primary cell cultures of exclusively epithelial cells which contain all cell types of the small intestine ^21^. We profiled transcriptional changes of intestinal organoids which were generated from young and aged mice of 2-3 and 20-22 months (respectively hereafter referred to as young and aged organoids) by bulk RNA-Seq. We sequenced six different organoid lines per age (in total 12 lines from different mice) in two separate experiments in which the organoids had been cultured for four or five weeks prior to sequencing (Fig 4a). This prolonged culture period ensured immune cell-free organoid cultures as confirmed by our bulk RNA-Seq results, in which we could not detect *Cd45*, a pan immune cell marker.

**Fig 4:**
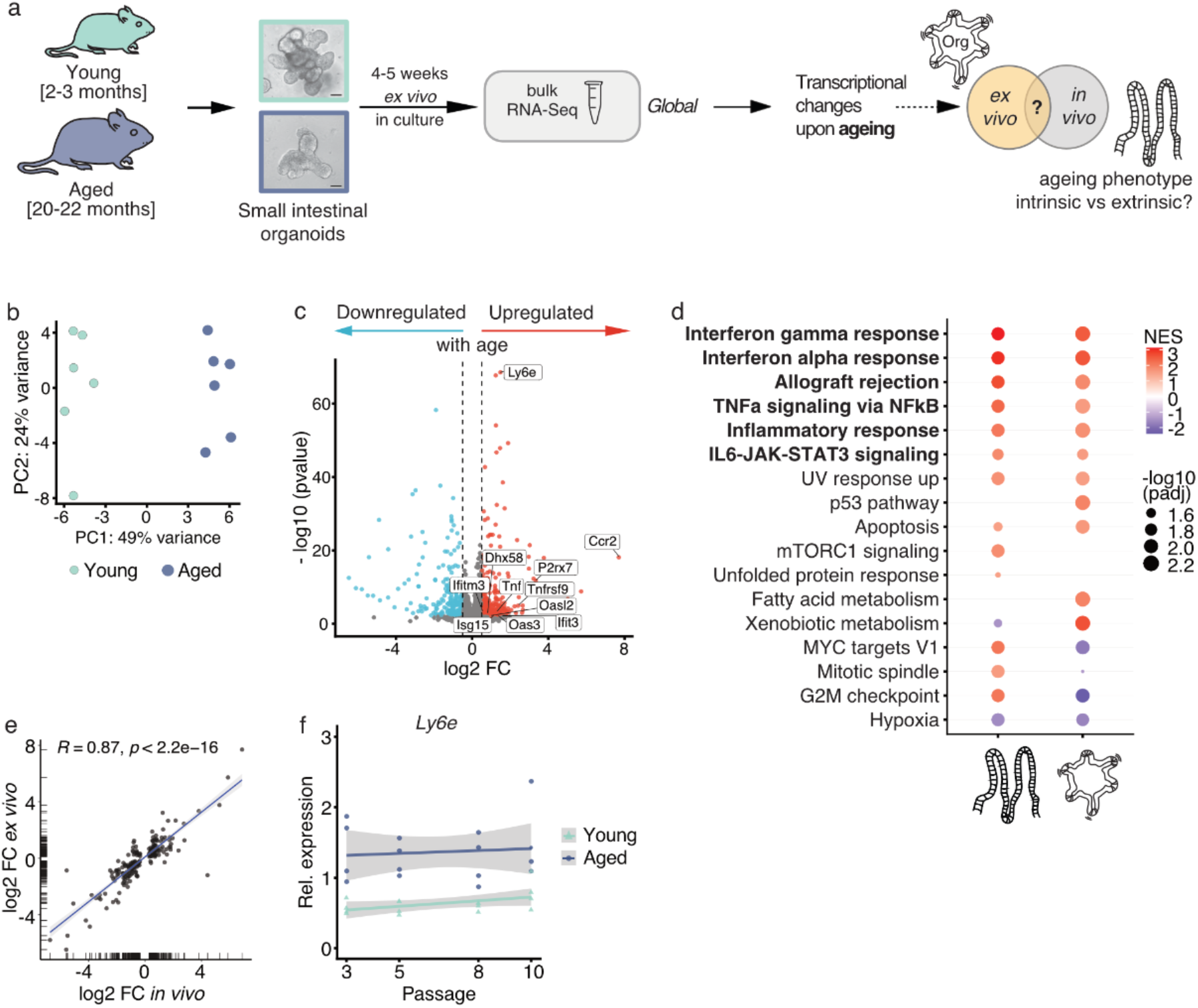
*Ex vivo* intestinal organoids reveal an epithelium-intrinsic inflammaging signature. **a)** Schematic of the experimental approach of bulk RNA-Seq of small intestinal organoids generated from young (2-3 months of age) and aged (20-22 months of age) mice and cultured for 4 or 5 weeks. **b)** PCA reveals the age of the mice (young: n=6, aged: n=6), from which the organoids were generated, as the first driving force to separate the samples based on transcriptional changes determined by bulk RNA-Seq. Variance explained by PC1 (x-axis) and PC2 (y-axis) is indicated in % on the respective axis. **c)** Organoids from aged mice show upregulation of immune-related genes. Volcano plot showing differentially expressed genes upon ageing (aged over young) in small intestinal organoids, red: significantly (FDR≤10%) upregulated (log2 FC > 0.5), blue: significantly (FDR≤10%) downregulated (log2 FC < (−0.5)), grey: not significantly changed and absolute log2 FC < 0.5. **d)** Gene set enrichment analysis indicates that the *in vivo* inflammaging phenotype is recapitulated in *ex vivo* cultured organoids. Side by side comparison of *in vivo* (left) and *ex vivo* (right) normalized enrichment score (NES) based on gene set enrichment analysis on hallmark gene sets (MSigDB); filtered for FDR≤5%, dot size indicates significance value (FDR), color code indicates NES. **e)** Correlation of the effects of ageing on the transcriptome as determined by bulk RNA-seq in intestinal organoids (y-axis) and *in vivo* intestinal epithelium (x-axis). A Deming regression line, as well as the Pearson correlation coefficient R and a p-value (p) of a test for correlation, are shown. Only genes significantly (FDR≤10%) affected in intestinal organoids are included. **f)** Expression of *Ly6e* is constant over the time period of ten passages (eleven weeks) in organoid cultures and higher in organoids derived from aged mice compared to young mice. Transcript levels were assessed in young and aged small intestinal organoids by qRT-PCR (normalized to *36B4*) (n=4 different organoid lines per age), data points representing the result of each biological replicate. (PCA: Principal component analysis, log2 FC: log_2_ fold change).

In a PCA, the age of the mice from which the organoids were generated, was the first driving force to separate the samples (Fig 4b), similar to what we had observed for freshly isolated epithelial cells. In fact, 49% of the variance between the samples could be explained by PC1, the age of the organoids. This clear separation of *ex vivo* cultures by age based on transcriptional changes indicated that age-related changes persist in culture for at least several weeks, independent of direct signals from the microenvironment. In total, we identified 599 genes significantly differentially expressed, from which 345 genes were upregulated and 254 genes were downregulated with age (Extended Data Fig 6a). Of these 598 DEGs, 147 genes overlapped with those detected *in vivo* (Extended Data Fig 6b), and 140 genes were regulated in the same direction (Extended Data Fig 6c). Within these 140 DEGs, we again identified genes involved in the immune response as upregulated upon ageing, such as *Ly6e, Ccr2, Ifitm3, P2rx7*, and *Tnf* (Fig 4c). Moreover, we observed a strong correlation between age-dependent DEGs detected in freshly isolated epithelial cells and in *ex vivo* cultured organoids (Fig 4e). In line, we identified a stable age effect across all samples of the same age when we analysed the *in vivo* and *ex vivo* bulk RNA-Seq data sets together (Extended Data Fig 6d, e).

To further assess the overlap of age-related transcriptional changes between intestinal organoids and *in vivo* intestinal epithelium from a functional perspective, we compared the enriched gene sets between both bulk RNA-Seq experiments. This analysis revealed enrichment for all six pathways of our previously defined inflammaging gene set in aged organoid cultures (Fig 4d), as observed in the freshly isolated intestinal epithelium. Besides the inflammaging gene set, also other gene sets showed overlapping enrichment between *in vivo* and *ex vivo* (UV response up and apoptosis) or a reduction (hypoxia). While other pathways, such as unfolded protein response or mTORC1 signaling, were only enriched *in vivo* but not changed in organoid cultures. Moreover, some gene sets (for example G2M checkpoint) showed an inversion from *in vivo* to *ex vivo*. Based on these results, we concluded that the inflammaging signature of the aged intestinal epithelium is partially maintained in culture. We wanted to test this hypothesis for an extended culture period to assess how long immune-related genes show higher expression levels in aged organoids compared to young ones. To this end, we performed a time-course experiment in which we cultured four young and four aged organoid lines over ten passages, corresponding to over two months in culture, and quantified gene expression by qRT-PCR. For *Ifitm3* we observed that the fold change of aged over young was varying but still positive after eight or ten passages (Extended Data Fig 6f). Furthermore, expression of *Ly6e* was consistently higher in organoids from aged mice compared to organoids from young mice independent of the time in culture, and also the absolute expression did not change over the analysed period of time (Fig 4f). These data further supported our hypothesis that aged *ex vivo* intestinal organoids acquire an inflammaging phenotype similar as observed *in vivo*, which persists over weeks in culture and appears to be stable towards changing conditions.

Many of the genes we found upregulated in the aged intestinal epithelium also show increased expression in IBD patients, including *Isg15, Ly6e, Stat1, Tnf, Oasl, Oas3, Ifit3*, and *Ifitm3* (Fig 1b, 4c) ^22,23^. We, therefore, tested if the inflammaging phenotype could be reduced by a pharmacological treatment tested for IBD ^24^. We treated organoids with TPCA-1 (Extended Data Fig 6g-j), a known IKKβ antagonist that primarily inhibits NF*κ*B signaling and with lower affinity STAT3 signaling ^25^. TPCA-1 has been shown to have anti-inflammatory effects in an IBD model by reducing the secretion of inflammatory cytokines, including Cxcl10 ^24^. We observed a robust downregulation of *Cxcl10* expression upon TPCA-1 treatment in young and aged organoids (Extended Data Fig 6g), confirming the efficient inhibitory function of TPCA-1. Next, we tested the potential of TPCA-1 to downregulate IBD-associated genes that were upregulated in the aged intestinal epithelium. *Isg15* and *Oasl2* were significantly downregulated by TPCA-1 in aged organoids to similar levels as observed in young organoids (Extended Data Fig 6h, i). However, we did not observe such an effect for *Ly6e*, which was not downregulated by TPCA-1 in young or aged organoids (Extended Data Fig 6j), indicating that several inflammation-associated signaling pathways act in concert during inflammaging, as suggested by GSEA.

In conclusion, here we identify that the inflammaging signature of the aged intestinal epithelium initially observed *in vivo* is retained *ex vivo* in intestinal organoids. Aged epithelial cells acquire an intrinsic inflammatory signature, which mimics a chronic inflammatory disease state and persists in culture in the absence of extrinsic signals.

### Epithelium-intrinsic inflammaging is associated with chromatin remodeling

Next, we aimed to identify how the epithelium-intrinsic inflammaging is maintained in culture without signals from the microenvironment. In the context of ageing, senescence-associated secretory phenotype (SASP) is a known potential source for cytokines that induce inflammatory responses in an autocrine and paracrine fashion, while interventions that eliminate senescent cells can ameliorate inflammation ^26^. To assess if SASP could promote the inflammation signature in organoid cultures, we measured cytokine levels (IL-6, IFNa, IP-10/Cxcl10, VEGF) in the medium of young and aged organoids by an electrochemiluminescence-based multiplex cytokine assay ^27^. With this approach, we were able to measure cytokines secreted by the organoids, however, we did not detect significant differences in cytokine levels between young and aged organoids (Extended Data Fig 6k-n). In line, also our bulk RNA-Seq data were not indicative for a senescence phenotype. Based on these results, we concluded that SASP is unlikely to fuel the epithelial inflammaging signature in aged organoids.

Previously a study on mouse tissue showed that ageing in the liver, heart and neural lineage was associated with an inflammatory response accompanied by epigenetic remodeling ^28^. Therefore, we were wondering whether epigenetic changes could underlie the inflammation-associated transcriptional changes observed in the aged small intestinal epithelium *in vivo* and explain how this transcriptional profile is maintained over weeks *ex vivo*. To test this hypothesis, we performed ATAC-Seq in young and aged small intestinal organoids that were cultured for three to five weeks prior to sequencing. The experiment aimed to map genome-wide open chromatin sites ^29^ and to assess if chromatin accessibility changes with age. The majority of the annotated peaks we identified were located as expected in open regions of the genome, mainly in distal intergenic (30%) and promoter regions (26%) (Extended Data Fig 7a). Remarkably, PCA showed a clear separation of young and aged samples based on their open chromatin sites (Fig 5a). 72% variation between the samples could be explained by PC1, which reflected the age of the organoids. This demonstrated that chromatin accessibility changes upon ageing in the intestinal epithelium and that many of these changes are maintained in culture for at least several weeks. Differential accessible region analysis revealed 2442 peaks that changed significantly upon ageing (Extended Data Fig 7b), from which 1291 indicated increased chromatin accessibility, typically associated with elevated transcription. For example, we detected increased chromatin accessibility by 6-fold for *Ccr2*, 5-fold for *Ly6e*, and 2.5-fold for *Dhx58* (Fig 5b, Extended Data Fig 7c, d). These genes are associated with immune response and inflammation and were also transcriptionally upregulated in aged organoids (Fig 4c). Next, we wanted to ascertain if changes in open chromatin sites could specifically underlie the epithelium-intrinsic inflammaging phenotype as observed on the transcriptional level in organoids. To this end, we plotted the significant DEGs of bulk RNA-Seq against the significant peaks of the ATAC-Seq data and highlighted all genes belonging to our inflammaging gene set (Extended Data Fig 7e). This analysis revealed that transcriptional changes upon ageing and differentially accessible regulatory regions correlated with each other and showed the same directionality in their regulation. For example, *Ccr2* showed increased chromatin accessibility in aged organoids, which coincided with transcriptional upregulation upon ageing. This correlation suggests that changes in transcript abundance upon ageing are partially predetermined by changes at the chromatin level and that these modulations are stable in culture for at least several weeks. Importantly, many of the genes that showed this trend belonged to our inflammaging gene set, including *Ly6e, Ccr2, P2rx7, Usp18, Il18r1*, and *Tnfrsf9*, supporting our hypothesis that epithelium-intrinsic inflammaging is enabled by initial chromatin remodeling.

**Fig 5:**
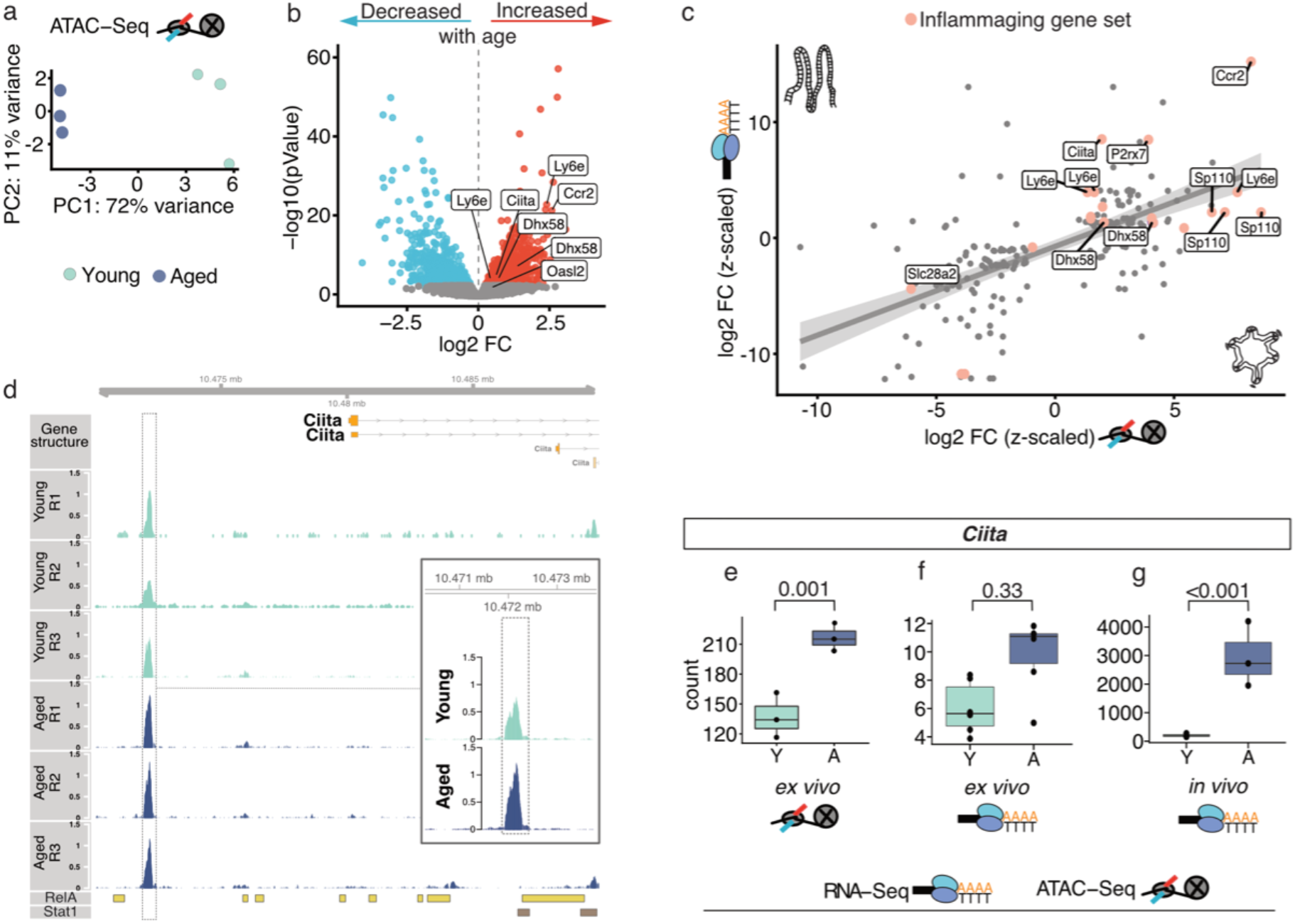
Epithelium-intrinsic inflammaging is associated with chromatin remodeling. **a)** Chromatin accessibility in young and aged organoids was determined by ATAC-Seq. PCA reveals the age of the mice (young: n=3, aged: n=3), from which the organoids were generated, as the first driving force to separate the samples. Variance explained by PC1 (x-axis) and PC2 (y-axis) is indicated in % on the respective axis. **b)** Chromatin accessibility in intestinal epithelial cells undergoes changes upon ageing and is persistent in *ex vivo* culture. Volcano plot showing significantly (FDR≤10%) changed peaks of chromatin accessibility in genetic regions/loci upon ageing (aged over young) in small intestinal organoids derived from young (n=3) and aged (n=3) mice; red: increased (log2 FC > 0), blue: reduced (log2 FC < 0), grey: not significantly changed. **c)** Scatterplot of the effects of ageing on the expression of genes as determined by bulk RNA-seq of *in vivo* intestinal epithelium (y-axis, z-scaled log2 FC) and on the chromatin accessibility as determined by ATAC-Seq in intestinal organoids (x-axis, z-scaled log2 FC). Genes of the inflammaging gene set are color-highlighted and a linear regression line indicates the agreement between the compared data sets (95 % CI shaded grey). Only genes significantly (FDR≤10%) affected *in vivo* intestinal epithelium are included. **d)** Regulatory region assigned to *Ciita* showed increased chromatin accessibility upon ageing in the small intestinal epithelium. Peak density is shown for all young (n=3) and aged (n=3) samples in genomic coordinates close to *Ciita* by R/Gviz. RelA and Stat1 binding sites determined by LOLA (Material and Methods) are indicated at the bottom. Inlet shows aggregated peak densities for all young and aged samples. Peaks of chromatin accessibility are color-coded by age of the sample. **e)** Count plots for *Ciita* for differentially open chromatin regions determined by ATAC-Seq of intestinal organoids and for transcript abundance as determined by bulk RNA-Seq in **f)** intestinal organoids and **g)** *in vivo* intestinal epithelium. Bars represent the average counts for all young and all aged samples, data points represent the counts of the biological replicates, adjusted p-Value is indicated on the respective comparison (log2 FC: log_2_ fold change).

To test if we can infer insights into the *in vivo* chromatin landscape from our organoid ATAC-Seq data, we performed the same analysis as before (Extended Data Fig 7e), while here we plotted the significant peaks of the organoid ATAC-Seq against the DEGs from the *in vivo* bulk-RNA Seq experiment (Fig 5c). We again observed a correlation between our bulk RNA-Seq and ATAC-Seq data, in particular for genes of the inflammaging signature (Fig 5c), suggesting that the elevated chromatin accessibility in the aged intestinal epithelium provides a permissive state for inflammatory responses, and this epigenetic potential is exploited *in vivo*.

Interestingly, we also detected a regulatory region with increased chromatin accessibility upon ageing for *Ciita* (Fig 5c - e, the transactivator of the MHC class II (MHC II) genes, which coincided with a 4-fold transcriptional upregulation *in vivo* (Fig 5c, g). In organoid cultures, *Ciita* and MHC class II genes are almost not expressed (Fig 5f), while co-culture with immune cells or direct treatment with IFN*γ* can restore their expression ^9^. Thus, organoids hardly express *Ciita* but retain the age-related increased DNA accessibility of the gene. This result indicates that *Ciita* displays a primed chromatin state for enhanced activation in the aged tissue, which stays unexploited in culture. Whereas *in vivo*, the interaction with the microenvironment, most likely the immune cells, might trigger the primed potential. This interaction could explain the marked transcriptional upregulation of *Ciita* and thus MHC class II genes in the aged *in vivo* intestinal epithelium.

To identify which external signal and transcription factors would be able to exploit the increased chromatin accessibility of *Ciita* and to facilitate the interpretation of our results of increased open chromatin states upon ageing in general, we made use of the R package LOLA (*Locus Overlap Analysis*). This package performs enrichment analysis for genomic loci overlaps ^30^, similar to a GSEA. In this analysis, significant overlaps between our ATAC-Seq data set (query regions of interest) and a reference database for differentially regulated genomic regions (CODEX and Encode) were assessed (Extended Data Fig 8a). With this analysis, we observed that the ATAC-Seq peaks of aged organoids overlapped with transcription factors binding sites associated with immune response and inflammation, such as Stat1, RelA, JunB, or Irf1. For *Ciita* we identified a RelA binding site in front of the chromatin region that showed increased accessibility in the aged epithelium (Fig 5d), indicating that NF*κ*B signaling, maybe in concert with Interferon signaling, is involved in the upregulation of *Ciita* and thus MHC class II genes in the aged intestine.

In conclusion, we show that aged intestinal organoids show a more open chromatin landscape in inflammation-associated genes, compared to young organoids. These changes are stable over several weeks in culture and indicate that the aged intestinal epithelium harbors an increased potential for inflammatory responses, which can be exploited differently *in vivo* and *ex vivo*.

### Levels of immune tolerance marker PD-1 and PD-L1 are elevated in the microenvironment of the aged intestine

Intestinal inflammatory states, as seen in IBD, are often associated with deregulated immune cell trafficking and altered immune cell activation ^31^. Part of the self-maintaining inflammatory process are MHC class II molecules, which are major sites of immune cell interaction by antigen presentation and thus display an important interface to immune cell communication ^32^.

Since we had observed in our *in vivo* bulk RNA-Seq data that MHC class II genes are highly upregulated in the aged epithelium (Fig 1d), we set out to assess the immunological status of the aged intestine on tissue level. To this end, we first aimed to confirm the findings of our transcriptomic data also on protein level by immunohistochemistry (IHC) stainings of small intestinal tissue sections (Fig 6a). We observed a marked increase of MHC class II protein abundance in the entire aged epithelium (Fig 6a, b). Exploring our *in vivo* single-cell RNA-Seq data, we identified a trend towards a more robust upregulation of MHC class II upon ageing in ISCs, TAs, enterocytes, and enterocyte progenitors, compared to secretory cell types (Fig 6c, Extended Data Fig 8b, c). These analyses show that the aged intestinal epithelium broadly upregulates MHC class II genes, which are critical communication sites with immune cells and suggest potential modulation of immune cells upon ageing. To assess the immunological homeostasis of the aged intestine, we analysed tissue sections of young and aged mice for helper (CD4^+^), cytotoxic (CD8^+^), and regulatory (FOXP3^+^) T cells (Tregs), as well as for the components that regulate and balance immune cell function between immuno-protection and immunopathology, PD1 and PD-L1 ^33^. We detected the great majority of immune cells in the intestinal mesenchyme of the villus region, while only a small proportion located at the level of crypts and rarely within the epithelium. (Fig 6d, g). For CD4^+^ T helper cells we observed a tendency of increased numbers, though not significant (Fig 6d, e), and we detected no changes for CD8^+^ cytotoxic T cells (Fig 6f). However, we found a significant increase of FOXP3^+^ Tregs (Fig 6g, h), which have a central role during immunological homeostasis ^34^. Moreover, we also detected increased levels of PD-1 (Fig 6i), which is transiently expressed in activated T cells and a marker for exhausted T cells, typically found during chronic infections ^33^. In line, we also found elevated levels of PD-L1 (Fig 6j), the PD-1 receptor that induces T cell exhaustion upon binding, further indicating deregulation of the immune equilibrium in the intestine upon ageing.

**Fig 6:**
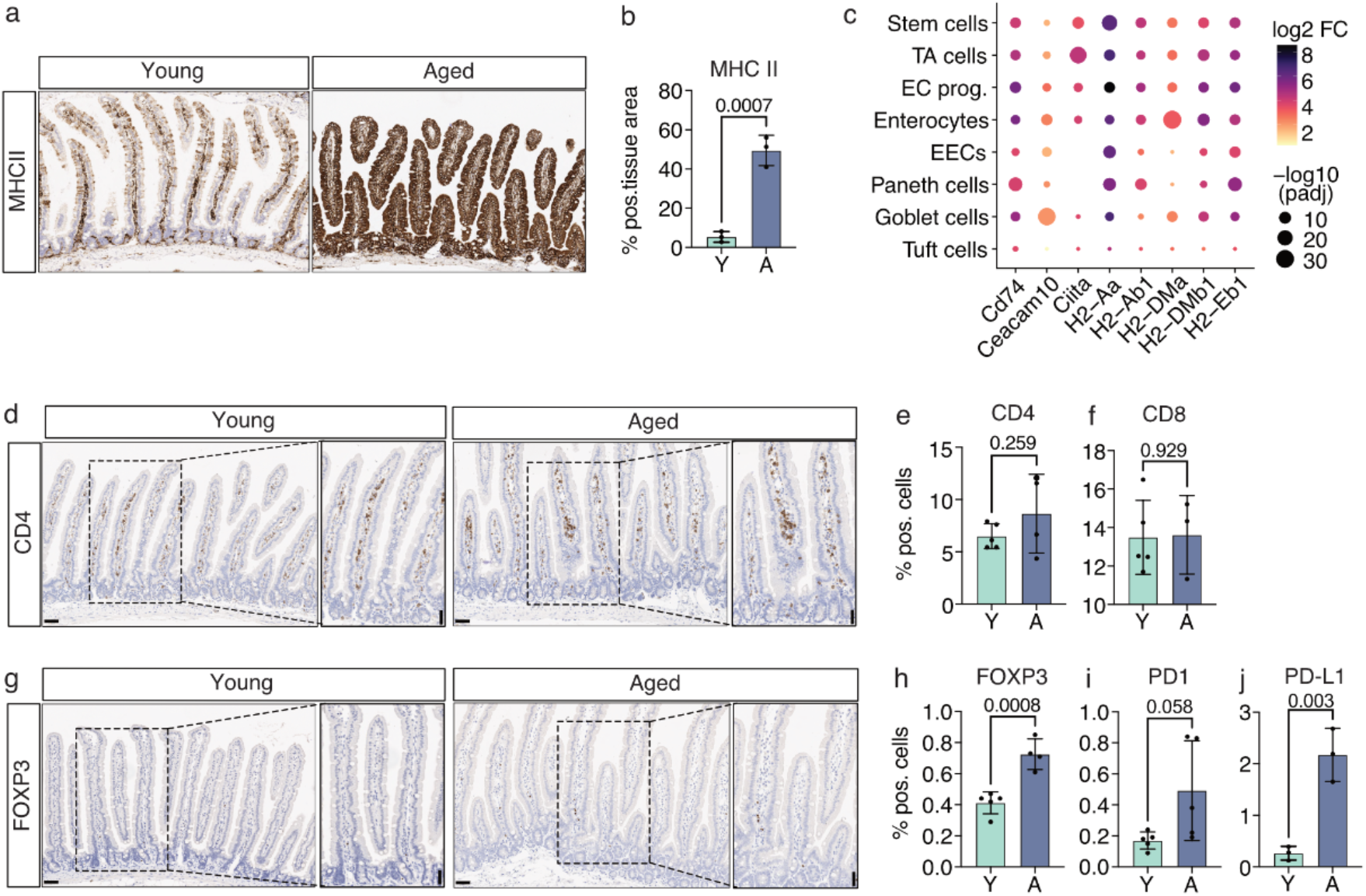
Levels of immune tolerance marker PD-1 and PD-L1 are elevated in the microenvironment of the aged intestine. **a, b)** Significant increase of MHC class II molecules on the aged intestinal epithelium as observed by **a)** IHC stainings for MHC class II molecules on whole intestinal tissue sections from young and aged mice (displayed are two representative images) and **b)** quantified as positive tissue area (n = 3 per young and aged). **c)** Dot plot showing upregulation of MHC class II molecules and related genes (x-axis) in the different cell types of the intestinal epithelium (y-axis) upon ageing. Dot size indicates significance, dot color indicates log 2 FC upon ageing (aged over young). **d, g)** IHC staining for **d**) CD4 and **g**) FOXP3 in whole intestinal tissue sections from young and aged mice, displayed images show representative stainings per age, the content of the dotted line frame is shown with higher magnification. **e-f)** + **h-j)** Quantification of IHC stainings as a percentage of positive cells for **e)** CD4, **f)** CD8, **h)** FOXP3, **i)** PD-1, **j)** PD-L1. Bars represent the average percentage for all young (n= 3-5) and all aged mice (n= 3-5), data points represent the percentage for the individual biological replicate, statistical significance was tested by an unpaired t-test (two-sided) and p-Value is indicated on the respective comparison. (MHC class II: Class II major histocompatibility complex molecules, IHC: Immunohistochemistry; scale bar = 50 μm, log2 FC: log_2_ fold change).

Thus, the aged intestine shows signs of elevated immune tolerance, including increased numbers of Tregs as well as upregulation of PD-1 and PD-L1. These data suggest that the chronically inflamed status of the intestinal epithelium is kept under control by enhanced immune tolerance, which could however in turn provide a ground for cancer initiation ^35^.

Taken together, our data demonstrate a cell-intrinsic inflammatory status of aged epithelial cells in the small intestine and a dysregulation of the immunological homeostasis. These findings indicate that the interaction of aged epithelial cells with their microenvironment, in particular immune cells, modulates immune cell function and homeostasis, an ageing phenotype known as immunosenescence. Our data provide evidence that epithelial cells, and thus non-hematopoietic cells, contribute to the systemic inflammation frequently observed upon ageing.

## Discussion

The intestinal epithelium forms a dynamic interface with its microenvironment, comprising immune cells, mesenchymal cells, and the microbiome. In this study, we asked the question, which ageing effects in epithelial cells of the intestine are epithelium-intrinsic and would persist in the absence of signals by the microenvironment. First, our study revealed an *in vivo* inflammaging phenotype in all epithelial cell types of the aged intestine, marked by MHC class II upregulation and most pronounced in enterocytes. Second, using *ex vivo* cultured intestinal organoids, we identified a cell-intrinsic inflammaging phenotype in aged epithelial cells that is reminiscent of inflammation signatures in human IBD patients and persisted over weeks in culture in the absence of signals from surrounding cells. This epithelium-intrinsic inflammaging was concurrent with chromatin remodeling at inflammation-associated loci, indicating that inflammaging is primed on the chromatin level. Within the intestinal microenvironment, we detected elevated levels of the immune tolerance marker PD-1 and PD-L1, indicating T cell exhaustion and dysregulation of the immunological homeostasis upon ageing.

Transcriptional profiling of freshly isolated *in vivo* intestinal epithelial cells first identified inflammaging as the most prominent ageing phenotype. Inflammaging is defined as low-grade chronic inflammation upon ageing, often accompanied by dysfunctional immunity, and contributing to frailty in the elderly in humans ^18,36^. Previous studies have described inflammation in the aged intestine, either on tissue level or in isolated crypts ^16,17^. The results of our bulk RNA-Seq experiment of *in vivo* sorted epithelial cells are in line with these previous findings and further allow us to allocate the inflammatory signature to epithelial cells. Moreover, we resolve the inflammaging signature at a single-cell level to assess which cell types are mostly affected. Besides inflammation, we also identified enrichment for mTORC1 signaling in the aged intestinal epithelium, which is also in agreement with recent studies ^14,37^.

Bulk RNA-Seq experiments do not distinguish between expression changes that are caused by changes in cell-type ratio or expression changes within individual cells and cell types. To resolve this question for our data set and to further assess which epithelial cell types are affected by the inflammaging phenotype, we profiled transcriptomes in single cells of freshly isolated intestinal epithelial cells of young and aged mice. We assigned all cells to the known epithelial cell types and compared the number of cells per cell type between young and aged samples. With this, we observed unaltered cell type ratios with age, indicating that the results of our bulk RNA-seq are mainly driven by transcriptional changes within individual cells and cell types upon ageing. Here, our results in part deviate from previous studies that investigated changes in secretory cell types along the proximal to distal axis of the small intestine. Two studies identified in the distal part of the aged small intestine increased numbers of Paneth cells ^14,38^, while another study reported a reduction of Paneth cells in the same region ^16^. Nalapareddy and colleagues showed increased numbers of Paneth cells per crypt and Goblet cells per villus in the aged epithelium of the proximal region ^12^. At the same time, a recent study observed the increase of Goblet cells only in the distal, not in the proximal part of the intestine ^17^. Thus, regional differences might not only underlie transcriptional changes in homeostasis ^17^, but also during ageing. Moreover, the differences in Paneth cell numbers in the proximal small intestine between our data and the observations by Nalapareddy and colleagues might be caused by different applied techniques and thus different sensitivity levels. Importantly, one main conclusion of our study is that cell type ratio changes do not primarily drive the observed inflammaging phenotype but instead expression changes within individual cell types. And this conclusion is further supported by the correlation between differentially expressed genes of our bulk RNA-Seq data and in the individual cell types of the single-cell approach (Extended Data Fig 5), where cell type ratio changes are fundamentally removed.

We further explored our single-cell RNA-Seq data to assess which cell types contribute to the inflammaging signature. Based on gene set enrichment analysis and calculation of an *inflammaging score* per cell type, we found that the inflammaging signature is present in all cell types and most pronounced in enterocytes. Moreover, we observed a robust increase of MHC class II genes on the transcript and protein level in aged intestinal epithelial cells (IECs). Upregulation of MHC class II genes has been described as a change upon ageing on the single-cell level also in several other mouse tissues and organs previously ^39^, implying MHC class II increase as a common ageing phenotype across hematopoietic and non-hematopoietic cell types. One function of MHC class II molecules is the antigen presentation to T cells. This interaction activates T cells, which subsequently upregulate PD-1 and binding to its ligand PD-L1 causes T cell exhaustion ^33^. We observed elevated levels of PD-1 and PD-L1 in the aged microenvironment of the intestine, which to our best knowledge has not yet been described before in the ageing intestine. This upregulation of PD-1 and PD-L1 as immune tolerance markers suggests a dysregulation of the immunological homeostasis and impaired T cell function upon ageing. A previous study demonstrated that intestinal epithelial cells, in particular Lgr5^+^ ISCs, function as non-conventional antigenpresenting cells by MHC class II ^9^. In our study, we did not assess if elevated levels of MHC class II molecules upon ageing also lead to increased antigen presentation to T cells. However, it is tempting to speculate if increased MHC class II levels in aged IECs directly cause elevated levels of exhausted T cells. In this case, aged epithelial cells would contribute to immunosenescence and systemic ageing.

Transcriptional alterations observed *in vivo* display the result of crosstalk between many different cell types, making it difficult to identify cell-intrinsic ageing effects. To assess to what extent the inflammaging signature of the intestinal epithelium is dependent on direct signals from the microenvironment, we exploited *ex vivo* organoid cultures that are devoid of non-epithelial cells. We observed a stable age effect that persisted for weeks in culture. Importantly, organoids showed an inflammaging phenotype in absence of surrounding immune cells, suggesting an epithelial immunity phenotype that is intrinsic to IECs. The functional role of non-hematopoietic cells in organ-specific immune responses has been highlighted in a recent study in 12 mouse organs ^40^ and together with our data, this further supports the idea that aged epithelial cells might have the potential to affect the functionality of other non-epithelial cells and propagate systemic ageing, for instance by immunosenescence.

A recent publication analysed the effect of culture on intestinal epithelial cells ^41^. In this study, the authors compared the transcriptome of freshly isolated crypts with that of organoids after two weeks of culture. In line with our results, they also observed an enrichment of immune response and antigen presentation signatures upon ageing in isolated crypts. Generally, they found that most ageing features were lost upon culture. Gene set enrichment analysis of our organoid RNA-Seq experiment showed that many but not all *in vivo* ageing signatures are recapitulated in culture, while importantly, we demonstrated that the inflammaging signature stably persisted in culture for over two months. Future work will be needed to further develop organoid cultures into more advanced co-cultures with for example immune cells or the microbiome, to fully recapitulate the ageing phenotype of interest.

Typically, inflammatory responses include the presence and action of immune cells, for example, to secrete cytokines and to trigger inflammatory signaling in surrounding cells. To address the question of how aged intestinal organoids maintain the inflammaging signature in absence of immune cells, we investigated open chromatin sites in our organoid cultures. The organisation of the epigenome and transcriptome undergoes substantial changes upon ageing, which was demonstrated in a set of murine organs, not including the small intestine ^28^. With our work, we show that the epithelium-intrinsic inflammaging signature coincides with increased chromatin accessibility in inflammation-associated loci. Future work will be required to disentangle what causes remodeling of open chromatin sites in these genomic regions and if, for example, recurrent infections over lifetime could enhance these modulations.

In summary, our study demonstrates that an inflammaging signature is present in all cell types of the aged intestinal epithelium and most pronounced in enterocytes. Intestinal organoid cultures reveal an epithelium-intrinsic inflammaging phenotype that persists in culture, which is supported by a more open chromatin landscape in inflammation-associated loci upon ageing. In the aged intestinal microenvironment, the immune tolerance markers PD-1 and PD-L show elevated levels, indicating T cell exhaustion and dysregulation of immunological homeostasis upon ageing. Our results provide evidence that epithelial cells might contribute to systemic inflammation and disbalance of immune homeostasis during ageing.

## Material and methods

### Mouse husbandry

In this study, male mice of the strain C57BL/6JRj were used. Young (2-3 months of age) and aged (20-22 months of age) mice were bought from Janvier Laboratories, where mice are maintained under specific-pathogen free conditions and housed in a controlled environment and health status. Animals are kept at a day/night cycle of 12h/12h, cages are equipped with enrichment and mice are fed with a standard diet of 21% protein ad libitum. Animals were sacrificed by cervical dislocation according to an institutional license DKFZ-223 and in agreement with the German federal and state regulations.

### Single-cell dissociation of small intestinal tissue and FACS sorting of epithelial cells

Proximal small intestinal tissue was isolated, cleaned and incubated for 5 min in ice-cold PBS with 10 mM EDTA (AM9261, Invitrogen), followed by gentle shaking for 1 minute (min). Tissue was transferred into a new PBS solution with 10 mM EDTA and 10 μM Y-27632 (Dihydrochloride Rock inhibitor, SEL-S1049, Biozol Diagnostica), and incubated for 30 mins at 4 °C; released epithelial cells were filtered through a 70 μm cell strainer (43-10070-40, pluriSelect). To obtain a single cell suspension of epithelial cells, cells were resuspended in Advanced DMEM/F12 (AD) (11540446, Gibco) supplemented with TrypLE (12604-013, Gibco) and DNase I (0.3 U/mL (07900, STEMCELL Technologies)) and incubated 45 mins at 37 °C and further filtered through a 20 μm cell strainer (43-10020-40, pluriSelect). Cells were centrifuged and resuspended in 600 μl Blocking buffer (PBS + 2 % BSA (B9000S, New England Biolabs)) and incubated for 5 min on ice. Cells were stained with anti-CD326 (EpCam)-PE conjugated (12-5791-81; Invitrogen) and anti-CD45-APC conjugated (17-0451082; eBioscience), both diluted 1:300 directly into the blocking buffer and incubated for 15 min on ice. Living cells were gated by DAPI dye exclusion, epithelial cells were isolated as EpCAM^+^, CD45^-^ cells and first 25 000 cells were sorted for singlecell RNA-Sequencing, and then 300 000 cells per sample for bulk RNA-Seq, into IntestiCult Organoid Growth Medium (06005; STEMCELL Technologies). Sorting was performed on a BD FACS Aria I cell sorter (100 μm nozzle size).

For bulk RNA-Seq, the cells were resuspended in 300 μl RLT buffer (79216; QIAGEN) with 1:100 ß-Mercaptoethanol (63689; Sigma Aldrich). RNA extraction was conducted via QIAGEN RNeasy Mini kit (74106; Qiagen), including an on-column DNase digestion using the RNase-free DNase Set (79254; Qiagen).

### Bulk RNA-Seq library generation & sequencing

Illumina sequencing libraries were prepared using the TruSeq Stranded mRNA Library Prep Kit (20020594, Illumina) according to the manufacturer’s protocol. Briefly, poly(A)+ RNA was purified from 500 ng of total RNA using oligo(dT) beads, fragmented to a median insert length of 155 bp and converted to cDNA. The ds cDNA fragments were then end-repaired, adenylated on the 3’ end, adapter ligated and amplified with 15 cycles of PCR. The libraries were quantified using Qubit ds DNA HS Assay kit (Life Technologies-Invitrogen) and validated on an Agilent 4200 TapeStation System (Agilent technologies). Based on Qubit quantification and sizing analysis, multiplexed sequencing libraries were normalized, pooled and sequenced on HiSeq 4000 single-read 50 bp with a final concentration of 250 pM (spiked with 1% PhiX control).

### Bulk RNA-Seq analysis of freshly isolated intestinal epithelium (1), intestinal organoids (2)

Raw sequencing reads were aligned using STAR aligner (vSTAR_2.6.1d) ^42^. To this end, the nf-core pipeline rnaseq (v1.4.2, doi: 10.5281/zenodo.1400710) was employed ^43^. A conventional DESeq2 (1.30.1) pipeline was used to perform differential expression analysis ^44^. (1) 23,703 or (2) 24,674 transcripts were detected. Variance stabilizing transformation was adopted to stabilize the variance of genes with low read count for visualization of the sample dispersion. Significantly differentially expressed genes were called within a threshold of FDR < 10 % and an absolute log2 FC > 0.5. Molecular signature and enrichment analysis were performed with the FGSEA package and hallmark signatures from MsigDB ^45,46^. Overlap analysis between experiment 1) and 2) was performed using the online Venn diagram tool (http://bioinformatics.psb.ugent.be/webtools/Venn/) by VIB / U Gent, Bioinformatics & Evolutionary Genomics.

### Single-cell RNA-Seq library generation & sequencing

Cells from the proximal small intestine were prepared for FACS sorting and living epithelial cells were gated by DAPI dye exclusion, epithelial cells were isolated as EpCAM^+^, CD45^-^ cells. Per samples, 25 000 cells were sorted into IntestiCult Organoid Growth Medium (0600-0, −2 and −3 combined, STEMCELL Technologies), centrifuged and directly processed for library generation with the 10x Genomics technology according to the manufacturer’s protocol. Library generation was performed using the Chromium Controller instrument, the Chromium Single Cell 3’ Library, and Gel Bead Kit v3, and the Chromium Single Cell B Chip Kit. All cleanup steps were conducted using AMPure XP beads (A6883, Beckman Coulter). In short, about 20,000 cells from each sample were loaded per lane of the chip. Gel Beads in Emulsion (GEMs) were successfully generated for all samples, reverse transcription of mRNA was followed by cDNA amplification using 11 or 12 PCR cycles. The final library was generated and amplified in a Sample Index PCR using 12 or 13 cycles. All libraries sequenced had an average fragment size of 483 - 515 bp. To minimize technical artifacts, we pooled the final libraries and sequenced the multiplex on 2 lanes of a NovaSeq 6000 instrument with 100 cycle S1 reagent kits (read1 (cell barcode + UMI): 28 bp, read2 (transcript): 94 bp, i7 (sample index): 8 bp).

### Single-cell RNA-Seq analysis

Raw, demultiplexed sequencing data was processed with ‘cellranger’ v3.0.1 ^47^ with parameters ‘--transcriptome=mm10-1.2.0’ and ‘--expect-cells=5000’. The filtered count matrices were further processed with Seurat v4.0.5: only cells with more than 10 000 UMI counts and with at most 15% of UMI counts from mitochondrial genes were selected for further analysis (Extended Data Fig 2 a). Cells from different replicates were integrated using Seurat’s ‘IntegrateData’ ^48^. For visualization, a tSNE embedding of cells was computed based on the first 10 Principal components of the integrated and scaled data, using a UMAP embedding of the same as initialization. Cells were assigned to their respective types based on the labels transferred from cells of ^20^ using Seurat’s ‘TransferData’ but refining class labels along the Stem-Enterocyte axis to obtain simpler cell types, less dependent on cell cycle phases. Absolute differences in cell type ratios between young and aged samples was tested for with a paired t-test. Fold changes of cell type proportions were tested for with DEseq2 with size factors based on total counts and a global overdispersion estimate. Original UMI counts from all cells of each gene, sample and - for cell type specific analyses - cell type were aggregated by summation to form pseudo bulk aggregates and differential expression testing was performed on them separately for each cell type using DEseq2, adjusting for Chromium run. PCA embedding of samples was computed using DEseq2’s ‘varianceStabilizingTransformation’ without adjusting for Chromium run. All single-cell RNA-seq analyses and visualizations have been performed in R v4.1.0 making heavy use of R packages ‘data.table’ and ggplot2 ^49^.

Fgsea ^45^ (the fast R-package implementation of the preranked gene set enrichment analysis algorithm (GSEA) ^50^) was used to perform the gene set enrichment analysis on each cell type based on a subset of the hallmark gene sets extracted from the Molecular Signature Database (MsigDB ^46^) collection. Those are gene sets that can be linked to the ageing process.

A smaller subset of 6 hallmark gene sets (HALLMARK_INTERFERON_ALPHA_RESPONSE, HALLMARK_INTERFERON_GAMMA_RESPONSE, HALLMARK_ALLOGRAFT_REJECTION, HALLMARK_TNFA_SIGNALING_VIA_NFKB, HALLMARK_INFLAMMATORY_RESPONSE, HALLMARK_IL6_JAK_STAT3_SIGNALING) was then combined to create the list of genes related to the inflammaging phenotype. That consisted of a total of 151 unique genes present in the single cell data. To associate an inflammaging score to each cell, later averaged to each cell type, the AddModuleScore function ^51^ from Seurat V4 ^52^ was used. The inflammaging score was then scaled to range from 0 to 1.

### Small intestinal organoid generation, maintenance & treatments

Small intestinal organoids were generated as described before ^21^. Proximal small intestine was isolated, cleaned and villi were removed gently with a coverslip. The intestine was cut into small pieces and applied to serial washing steps in pre-chilled PBS and Penicillin-Streptomycin (100U) (15140122, Life Technologies). For crypt isolation the intestinal tissue pieces were transferred into 10 ml of Gentle Cell Dissociation Reagent (07174, STEMCELL Technologies) and incubated for 15 mins at room temperature (RT). After incubation, the tube was shaken vigorously for approximately to release the crypts and the solution was passed through a 70 μm cell strainer (43-10070-40, pluriSelect) into 30 ml PBS (20012027, Gibco) to stop the enzymatic digest and to exclude larger tissue pieces. Crypts were resuspended in Matrigel (Growth Factor Reduced, 356231, Corning) to obtain a final crypt concentration of 10 000 crypts per ml, which corresponds to 200 crypts per well of a 48-well plate. 20 μl of Matrigel-Crypt-Mix per well were seeded in a 48-well plate and 250 μl of IntestiCult Organoid Growth Medium (06005; STEMCELL Technologies) with Primocin (1:500, always added fresh) (ant-pm-05, InvivoGen) were added to each well and incubate at 37°C, in a humidified CO2 incubator. Medium was renewed every 2-3 days and organoids were split every 7 days. For TPCA-1 treatment, organoids were treated with 5μM TPCA-1 (T1452, Sigma Aldrich) and controls with DMSO at the same final concentration for 24 hrs.

### Organoid RNA isolation, cDNA generation and qRT-PCR

Organoids from typically 3 wells were harvested in 300 μl Cell Recovery Solution (354253; Gibco) and lysed in 300 μl RLT lysis buffer. Samples were filtered with QIAShredder (79656; Qiagen) columns, followed by RNA isolation with QIAGEN RNeasy Mini kit (74106; Qiagen), including an on-column DNase digestion using the RNase-free DNase Set (79254; Qiagen). cDNA was prepared from 0.5-1 μg total RNA with the QuantiTect reverse transcription kit (205311; Qiagen), according to the manufacturer’s protocol. cDNA was diluted to 10 ng/μl and 20 ng were used for each qRT-PCR reaction, using Maxima SYBR Green (K0221; Life Technologies) on the Lightcycler480 (Roche) in a 384-well format. Relative expression of each gene was calculated relative to the housekeeping gene *36B4* as 2^-Δ*C*t^. Oligonucleotide sequences are listed in Supplementary Table 1, all primers were validated prior to use in a cDNA dilution series.

### Organoid single-cell dissociation and ATAC-Seq library generation

Per sample, organoids from 8 wells (48 well-plate) were harvested for single cell dissociation. Matrigel was dissolved using ice-cold Advanced DMEM/F12 (AD) (11540446, Gibco), washed once with AD and then subjected to an enzymatic digest with TrypLE (12604-013, Gibco) + DNase I (07900, STEMCELL Technologies) at a final concentration of 0.1 mg/ml for 5 mins at 37 C. Digestion was stopped by addition of AD + 10% FBS. Cells were washed twice with AD and then passed through a 20 μm cell strainer (43-10020-40, pluriSelect) (pre-equilibrated with AD). ATAC-Seq library generation was based on Buenrostro et al., 2015 ^29^. Organoids were dissociated to single cells as described before. 100 000 single cells were treated with 50 μl cold lysis buffer (0,1% NP-40 (492016, Millipore), 0,1% Tween-20 (P1379, Sigma Aldrich), 0,01% Digitonin (G9441, Promega) in Resuspension buffer (RB)) for 3 min on ice. Then 1 ml Washing buffer (0,1% Tween-20 in RB) was added to the cells and centrifuged for 10 min at 4C at 500g. Supernatant was discarded and the pellet including the nuclei was resuspended in a 50 μl transposition reaction mix (Illumina Tagment DNA Enzyme and Buffer Kit, 20034210).

Transposition reaction and PCR amplification was carried out according to Buenrostro et al., 2015, using Nextera i5 and i7 index adapters to generate double index libraries. Libraries were purified via double-sided bead purification using AMPure XP beads (A6883, Beckman Coulter) and the library quality and quantity were assessed using Agilent High Sensitivity DNA Bioanalysis Chips (5067-4626, Agilent) and QuBit, respectively.

### ATAC-Seq sequencing & analysis

ATAC-Seq libraries of all samples were pooled and sequenced on a NextSeq, Paired-end 75bp, this generated a total of 727 107 568 reads of which 725 802 530 passed quality control. Raw reads were processed using the end-to-end integrated nf-core pipeline *atacseq* (version 1.2.1, doi: 10.5281/zenodo.2634132, ^43^). The results of the *broadPeak* analysis were used to pursue downstream analysis. The analysis was based on the ENSEMBL mouse genome release GRCm38.p6. Given the insights of the comparative analysis from Gontarz and colleagues we analysed our data set accordingly employing R/edgeR to call differentially open chromatin regions ^53,54^. As statistic for gene set enrichment analysis, we employed the equation stat= qnorm(1-pvalue/2)*sign(log2FC). Resulting differentially open peaks were annotated using ChipSeeker ^55^ and the M. musculus mm10 UCSC transcript database (TxDb.Mmusculus.UCSC.mm10.knownGene). For PCA analysis data was variance stabilized using DESeq2. Differentially open peaks were called significant if their Benjamini Hochberg corrected p-value was below 0.1 (10 % FDR). For comparing ATAC-Seq differentially open chromatin regions with differentially expressed transcripts, results were merged by the official gene symbol. The log_2_ fold changes were z-scaled and results plotted. A linear regression shows the correlation of the two distributions. In order to understand the transcription factors involved in the regulation of transcripts by open chromatin regions, we employed Locus Overlap Analysis (LOLA, ^30^) using the LOLACore database and differentially open chromatin regions in genes assigned to the HALLMARK of interferon gamma response ^46^. Results were visualized using R/Gviz and R/GenomicRanges ^56,57^.

### Cytokine assay

For the cytokine assay, we used the electrochemiluminescence-based multiplex assay ^27^ (K15069L-1; MSD U-PLEX Biomarker Group 1 (Mouse) Multiplex Assay; Meso Scale Discovery, Inc.) according to manufacturer’s protocol. We measured the medium that had been added to organoid culture for 2 days before measurement. 4 biological replicates per age were measured in technical duplicates.

### Histology, immunohistochemistry and quantification

Proximal small intestinal pieces were cleaned and fixed as swiss roll in 4% paraformaldehyde overnight at RT, paraffin embedded and sectioned (2 μm). Immunohistochemistry was performed on a Bond max System (DS9800; Leica Biosystems), using the Bond Polymer Refine Detection Kit (DS9800; Leica Biosystems). Sections were first applied to a peroxide blocking step for epitope retrieval using Epitope Retrieval Solution I and II (AR9640, Leica Biosystems), followed by primary antibody incubation for 30 mins at RT (PD-L1: 1 hr incubation time) and secondary antibody for 20 mins at RT used antibodies are listed in Supplementary Table 2. DAB enzymatic staining development was performed for 10 mins at 37C, using the Enzyme Pretreatment Kit (AR9551; Leica Biosystems) and followed by a counterstaining with Hematoxylin (AR9551; Leica Biosystems). Slides were scanned with a Leica Aperio AT2. For quantification the QuPath analysis software ^58^ was used. In brief, regions for measurement were selected, followed by the positive cell detection command, which counts all cells and all DAB-positive cells, resulting in the percentage of positive cells in the region of interest. Intensity threshold parameters for the positive cell detection command were as follows: Score compartment: Nucleus: DAB OD mean; Threshold: 0.15, 0.2 or 0.3 depending on the analysed staining while thresholds for the same staining were not changed between samples. MHC class II staining was analysed with the Fiji software ^59^ using the threshold command first to select tissue area and second to detect DAB-positive tissue areas. Statistical significance was tested by an unpaired t-test (two-sided). 3-4 biological replicates per age were stained and quantified.

### Statistics and Reproducibility

All statistical tests were performed with the GraphPad Prism Software (Version 9.2.0) or in the R program. All experiments include at least 3 biological replicates (biological replicate is defined as individual animal or organoid line derived from an individual animal) per age group or per treatment. Statistical details are described in the respective figure legend and exact p-values are plotted within the respective graph.

## Code availability

All source code needed to reproduce the figures as well as figure source data is available at: https://github.com/boutroslab/Supp_Funk_2021

## Data availability

All sequencing data that are presented in this study will be available upon publication at Gene Expression Omnibus under the accession ID GSE190286.

## Author contributions

MCF and MB conceived the study. MCF, ET and ZA performed experiments. JGG analysed single-cell RNA-Seq data, FH and MCF analysed bulk RNA-Seq. FH performed ATAC-Seq data analysis and EV performed further bioinformatic analyses. JH, DH performed IHC stainings. MCF, OS, MH and MB supervised the study; and MCF wrote the manuscript with contributions by MB. All authors edited and agreed on the manuscript.

## Acknowledgements

We are grateful to Kim E. Boonekamp, Fillip Port, Christian Scheeder and Josephine Bageritz, and the rest of the Boutros lab for valuable discussions. We thank Katharina Bauer and Jan-Philipp Mallm from the DKFZ single-cell open lab and DKFZ Flow Cytometry Core Facility for support with single-cell RNA-Seq experiments. The DKFZ Genomics and Proteomics Core Facility for support in sequencing bulk RNA-Seq. We further thank Stefania Del Prete and Duncan T. Odom for their support with the ATAC-Seq experiment and David Ibberson for help with sequencing the ATAC-Seq libraries. This work was supported by the Helmholtz Alliance “Aging and Metabolic Programming, AMPro” (MCF, MB). Work in the laboratory of MB & OS was supported, in part, by an ERC-Synergy Grant (“DECODE”) from the European Research Council (ERC).

---

**Supplementary Table 1.**
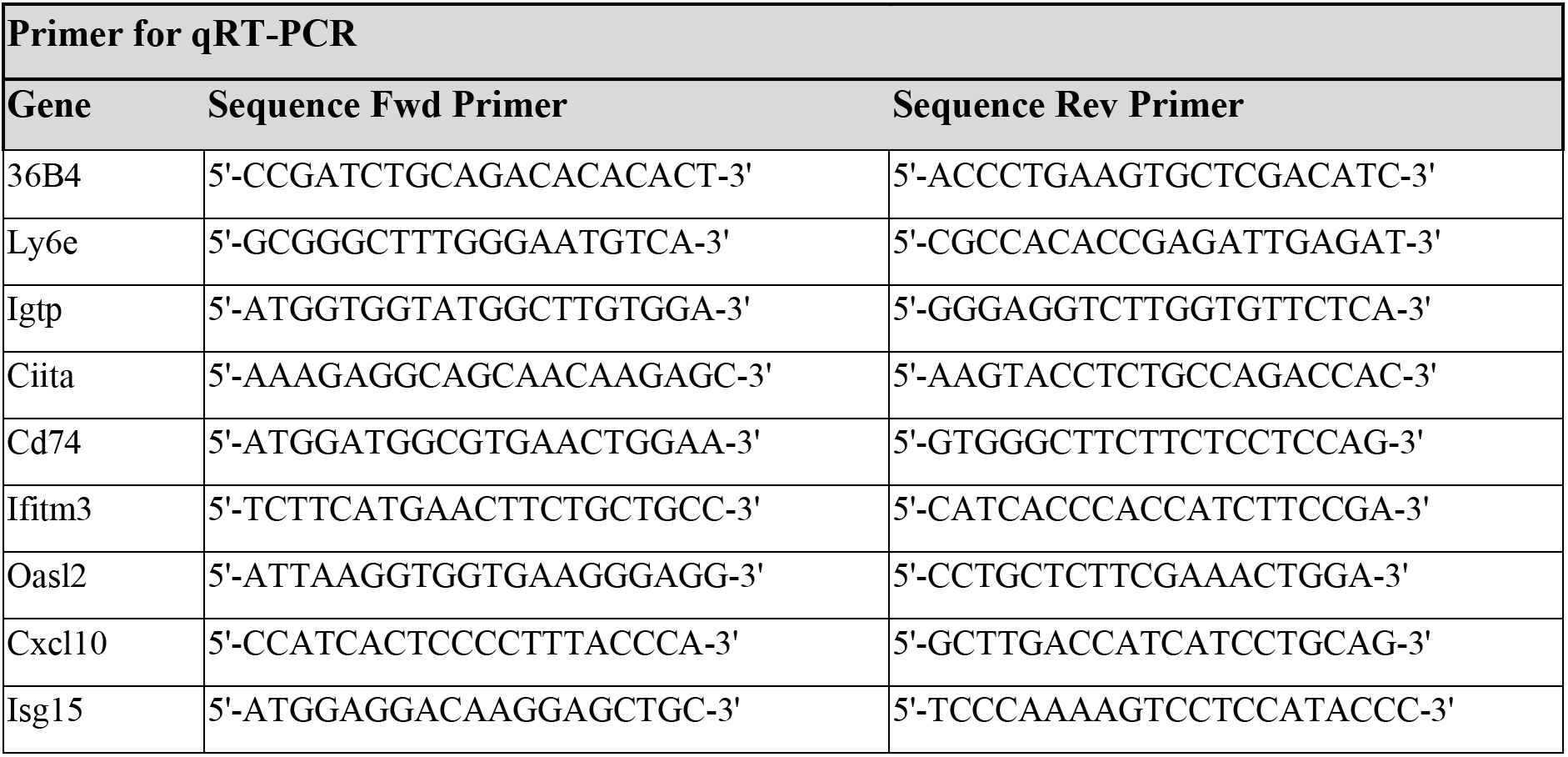

**Supplementary Table 2.**

| **Antibodies for immunohistochemistry** |  |  |  |
| --- | --- | --- | --- |
| **Primary Antibody** | **Company, Catalog number** | **Species** | **Dilution** |
| CD4 | eBioscience, 14-9766 | rat | 1:1000 |
| CD8 alpha | Invitrogen, 14-0808-82 | rat | 1:200 |
| FOXP3 | Cell Signalling, 98377S | rabbit | 1:50 |
| PD-1 | R&D, AF1021 | goat | 1:200 |
| PD-L1 | Cell Signalling, 649888 | rabbit | 1:50 |
| MHC class II | Novus Biologicals, NBP1-43312 | rat | 1:500 |
| **Secondary Antibody** | **Company, Catalog number** | **Species** | **Dilution** |
| anti-rat-HRP | Jackson Immuno Research, 312-005-045 | rabbit | 1:1000 |
| anti-goat-HRP | DAKO, P0449 | rabbit | 1:300 |
| anti-rabbit-HRP | Leica, DS9800 (Bond Polymer Detection Kit) | Antibody is part of the kit DS9800 and all reagents are provided ready to use |  |

## Extended Data Figures

**Extended Data Figure 1:**
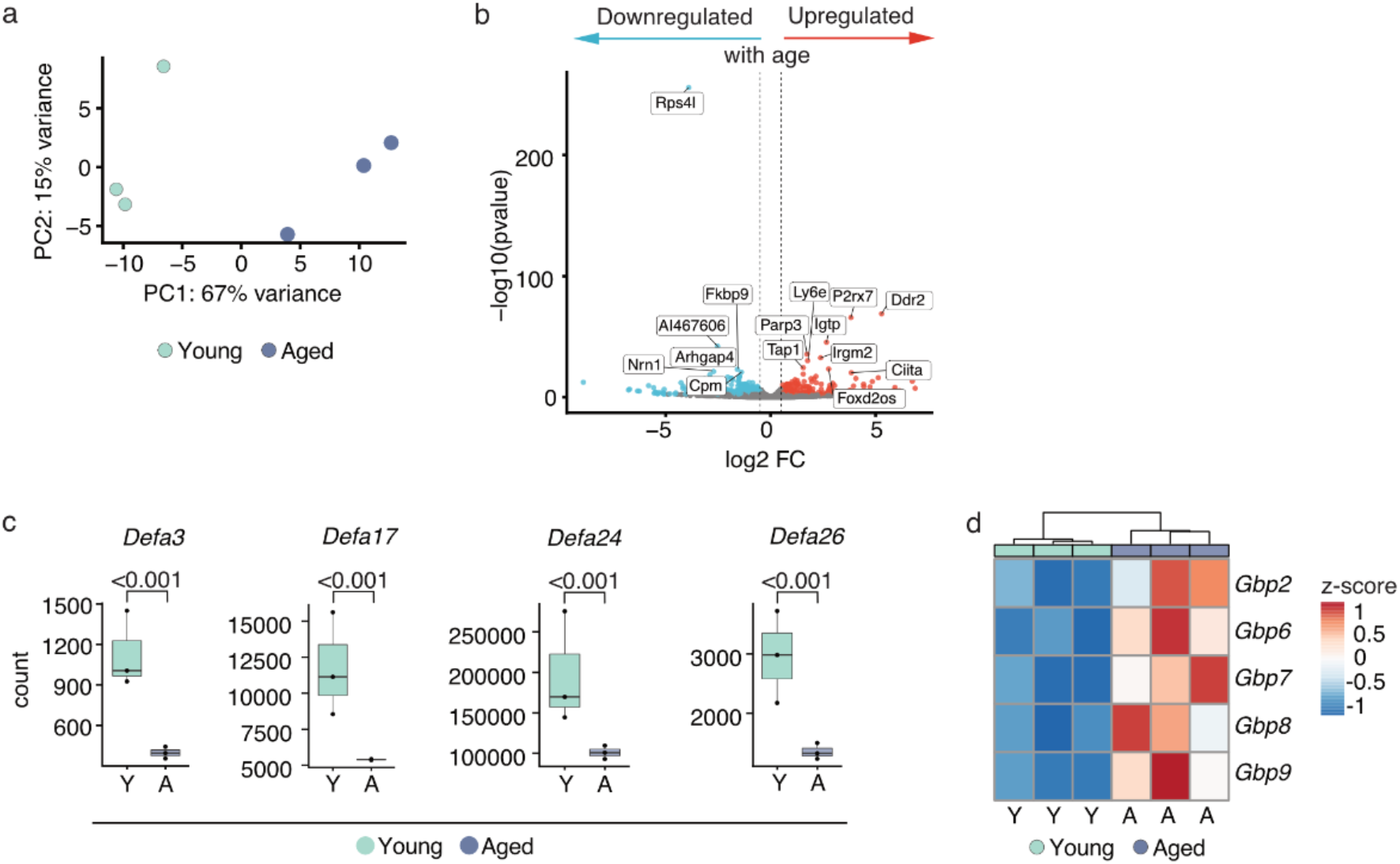
Transcriptional changes in the aged intestinal epithelium by bulk RNA-Seq. **a)** Age of the mice is the first driving force to separate the samples of the bulk RNA-Seq experiment, as shown by Principal component (PC) analysis of freshly isolated small intestinal epithelial cells of young (n=3) and aged (n=3) mice. PC1 and PC2 are plotted on the x- and y-axis, variance explained by PC1 and PC2 in %. **b)** Volcano plot depicting differentially expressed genes upon ageing (aged over young) in the small intestinal epithelium, red: significantly (FDR≤10%) upregulated (log2 FC > 0.5), blue: significantly (FDR≤10%) downregulated (log2 FC < (−0.5)), grey: not significant and absolute log2 FC < 0.5; gene names of the 15 most significantly changed genes upon ageing are plotted, **c)** Significant downregulation of alpha Defensins upon ageing. Count plots based on *in vivo* bulk RNA-Seq experiment for *Defa3, Defa17*, *Defa24*, and *Defa26*. Bars represent the average counts for all young (n=3) and all aged mice (n=3), dots represent the counts of the individual sample, adjusted p-Value is indicated on respective comparison, **d)** Upregulation of genes encoding for guanylate-binding proteins (*Gbp*) upon ageing in the small intestinal epithelium. Relative mean expression as z-score per *Gbp* gene (row) and sample (column) (3 young, indicated by “Y”, 3 aged, indicated by “A”), color code indicates z-score (log2 FC: log_2_ fold change).

**Extended Data Figure 2:**
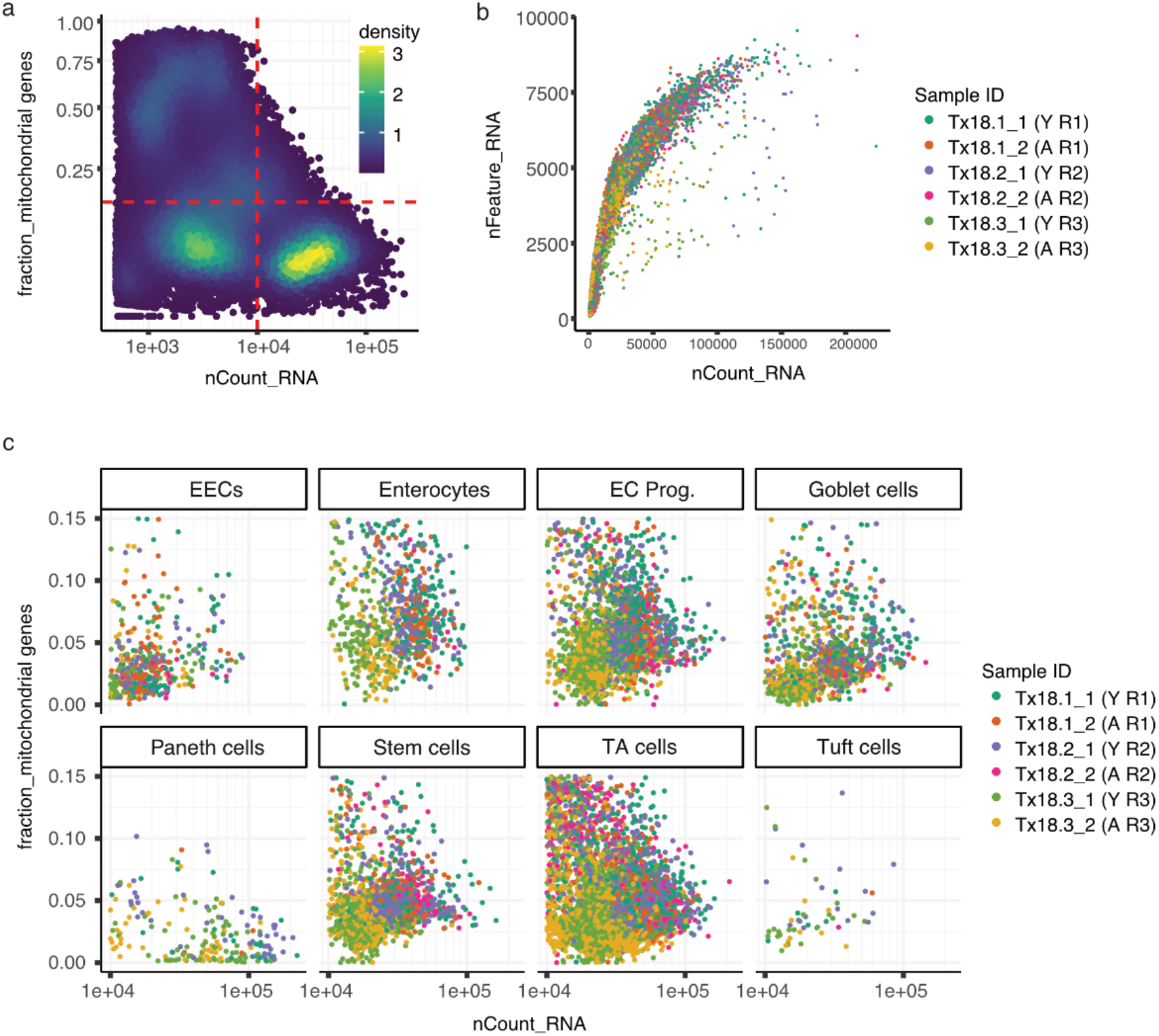
Quality control of single-cell RNA-Seq from small intestinal epithelium. **a)** Fraction of UMI counts from mitochondrial genes and total UMI counts of all droplets that passed cellranger’s filters. To restrict the analysis to alive and intact cells, transcriptomes with more than 10 000 total counts (dashed vertical red line) and less than 15% of counts from mitochondrial genes (dashed horizontal red line) were selected, **b)** The number of genes observed over the total number of UMI counts for each cell shows the typical saturating relationship for each sample (indicated by color), **c)** Quality control plot similar to **a)** but subset to the selected high quality cells as defined in **a)** and split up by the assigned cell type and colored by sample indicates no strong bias against certain cell types due to the QC filtering (EECs: enteroendocrine cells, EC prog.: enterocyte progenitors, TA cells: transit-amplifying cells).

**Extended Data Figure 3:**
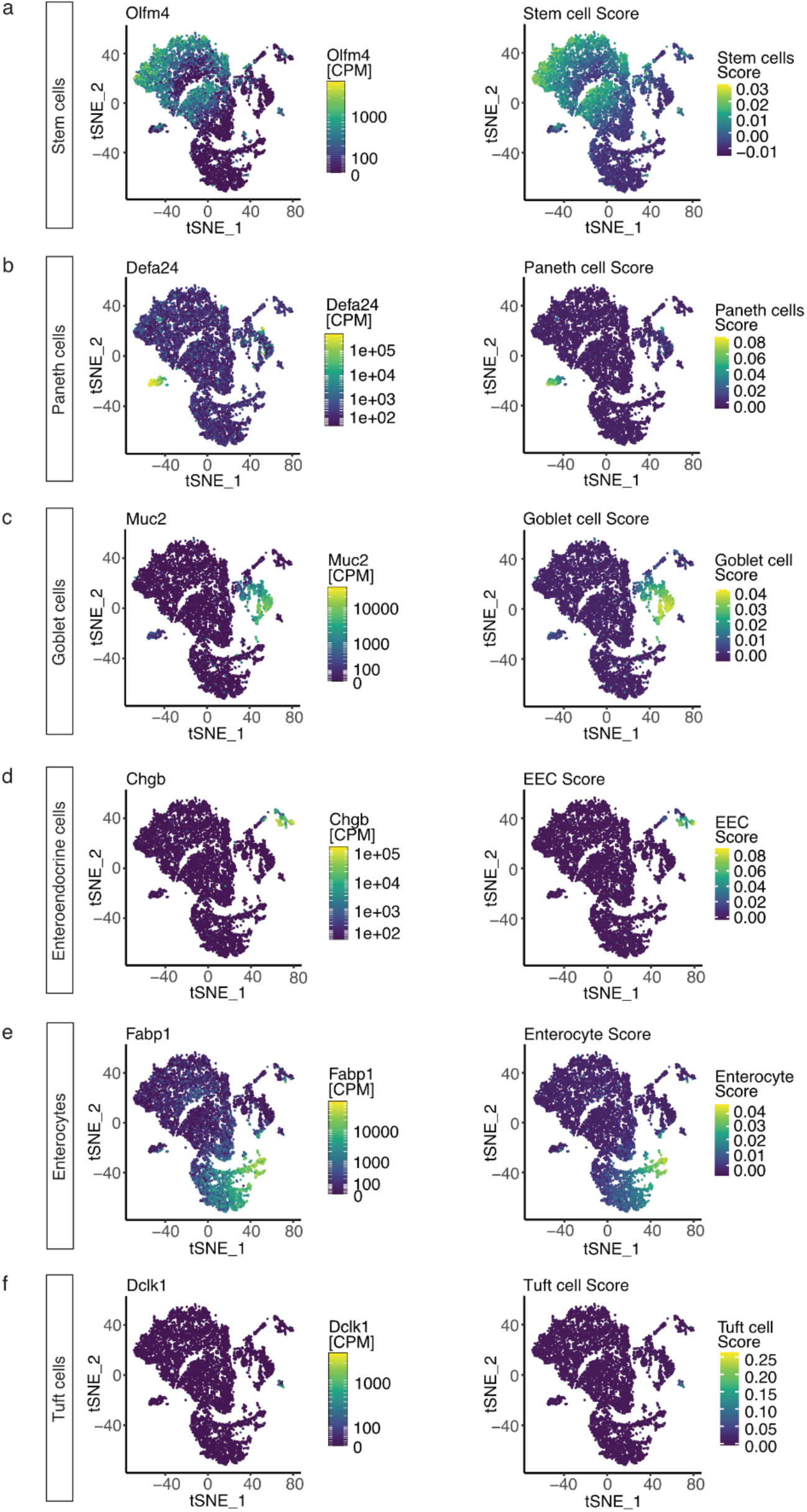
Small intestinal cell type markers and score of *in vivo* single-cell RNA-Seq. Cell type assignment was performed based on known cell-type marker sets (see Material and Methods). tSNE visualization of all sequenced cells (13 360) and color-coded for expression of one representative cell type marker per cell type (expression in CPM) and for cell type score, based on the cell type marker set. **a)** Stem cells marker *Olfm4* and Stem cell score, **b)** Paneth cell marker *Defa24* and Paneth cell score, **c)** Goblet cell marker *Muc2* and Goblet cell score, **d)** Enteroendocrine marker *Chgb* and Enteroendocrine cell score, **e)** Enterocyte marker *Fabp1* and Enterocyte score, **f)** Tuft cell marker *Dclk1* and Tuft cell score (CPM: counts per million mapped reads, EEC: Enteroendocrine cell).

**Extended Data Figure 4:**
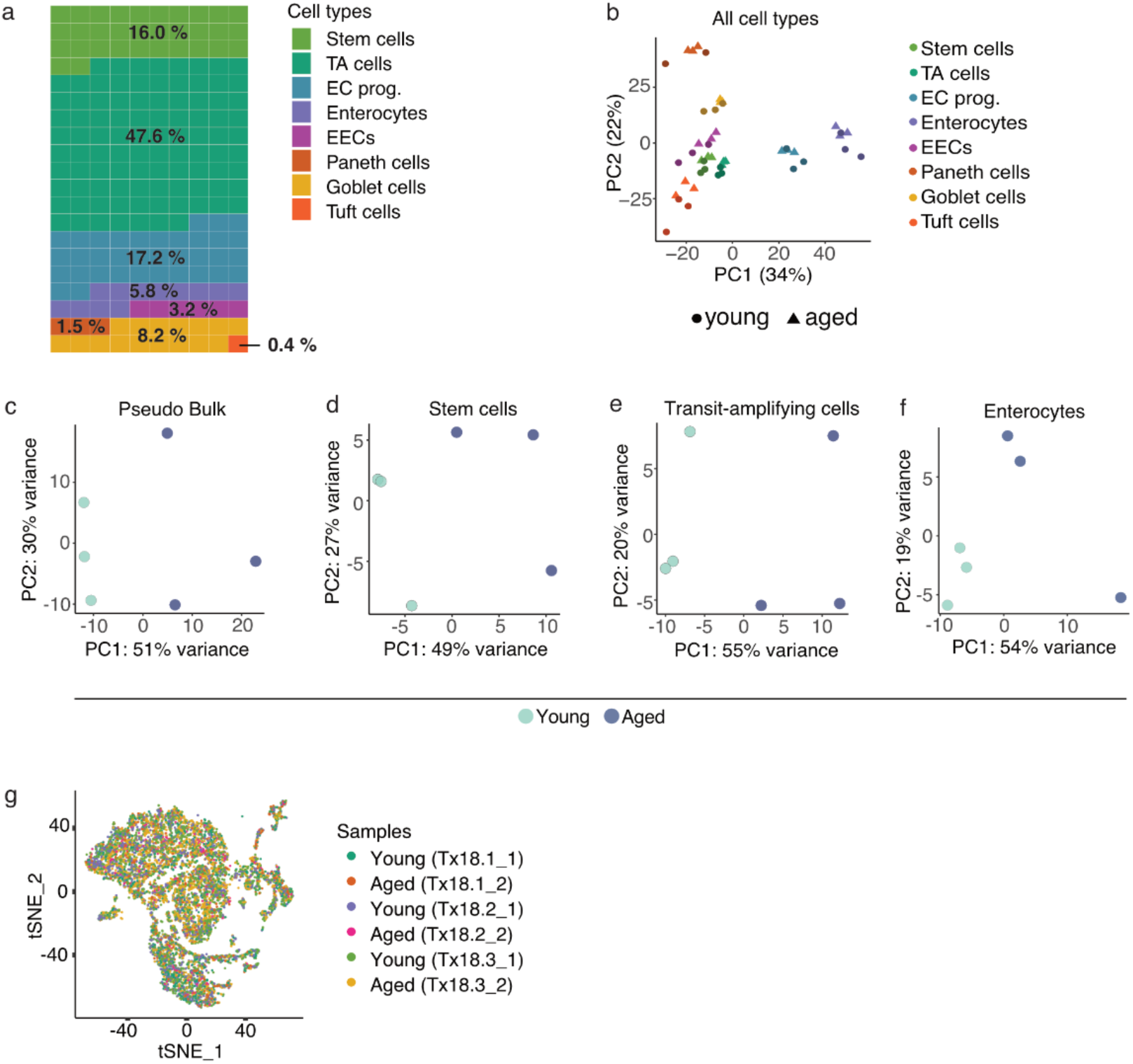
Single-cell RNA-Seq of young and aged small intestinal epithelium. **a)** Cell type proportions across all samples in %, represented in a waffle plot. Each square of the waffle plot represents 0.5% of all cells, **b)** PCA plot for all samples (young and aged) and all cell types per sample. Young and aged samples cluster primarily according to their cell type identity, **c)** PCA plot for pseudo bulk, samples separated by age along PC1 with a variance of 51%. **d)** PCA plot for stem cells, samples separated by age along PC1 with a variance of 49%. **e)** PCA plot for transit-amplifying cells, samples separated by age along PC1 with a variance of 55%. **f)** PCA plot for Goblet cells, samples separated by age along PC1 with a variance of 54%. **g)** tSNE visualization of all sequenced cells (13 360) and color-coded by the sample they originated from (TA: transit-amplifying, EC prog.: enterocyte progenitors, EECs: enteroendocrine cells).

**Extended Data Figure 5:**
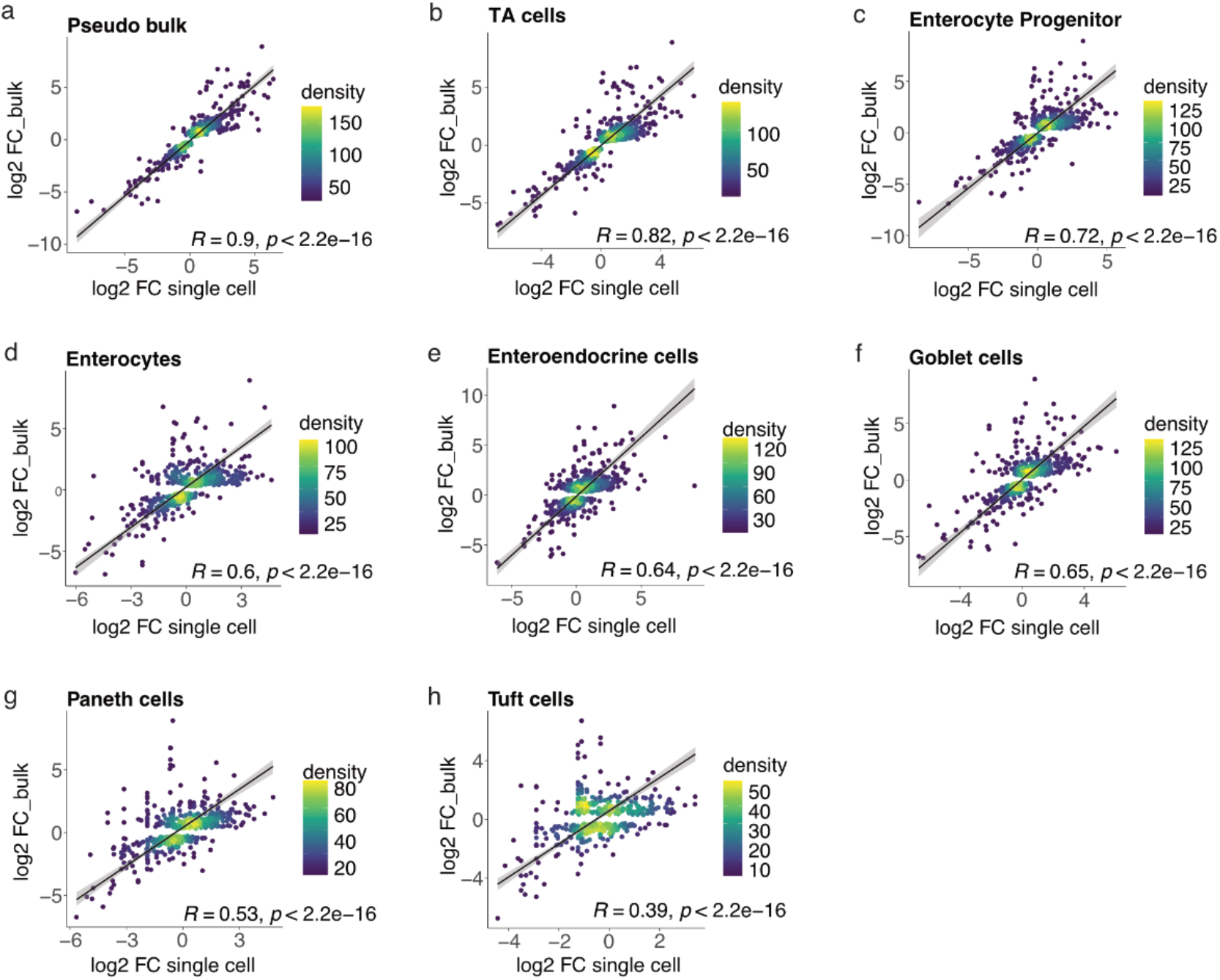
Effects of ageing on the transcriptome observed in bulk tissue correlate with the individual cell types of the intestinal epithelium to varying degrees. Correlation of the effects of ageing on the expression of genes in the small intestinal epithelium as determined by bulk RNA-seq (y-axis) and by single-cell RNA-seq (x-axis) for a pseudo bulk aggregate **(a)** as well as for the individual cell types **(b-h)**. Genes are colored by the density of points in the respective plot area. A Deming regression line as well as the Pearson correlation coefficient (R) and a p-value (p) of a test for correlation are shown in black. Only genes significantly (FDR≤10%) affected in bulk are included. (TA: transit-amplifying).

**Extended Data Figure 6:**
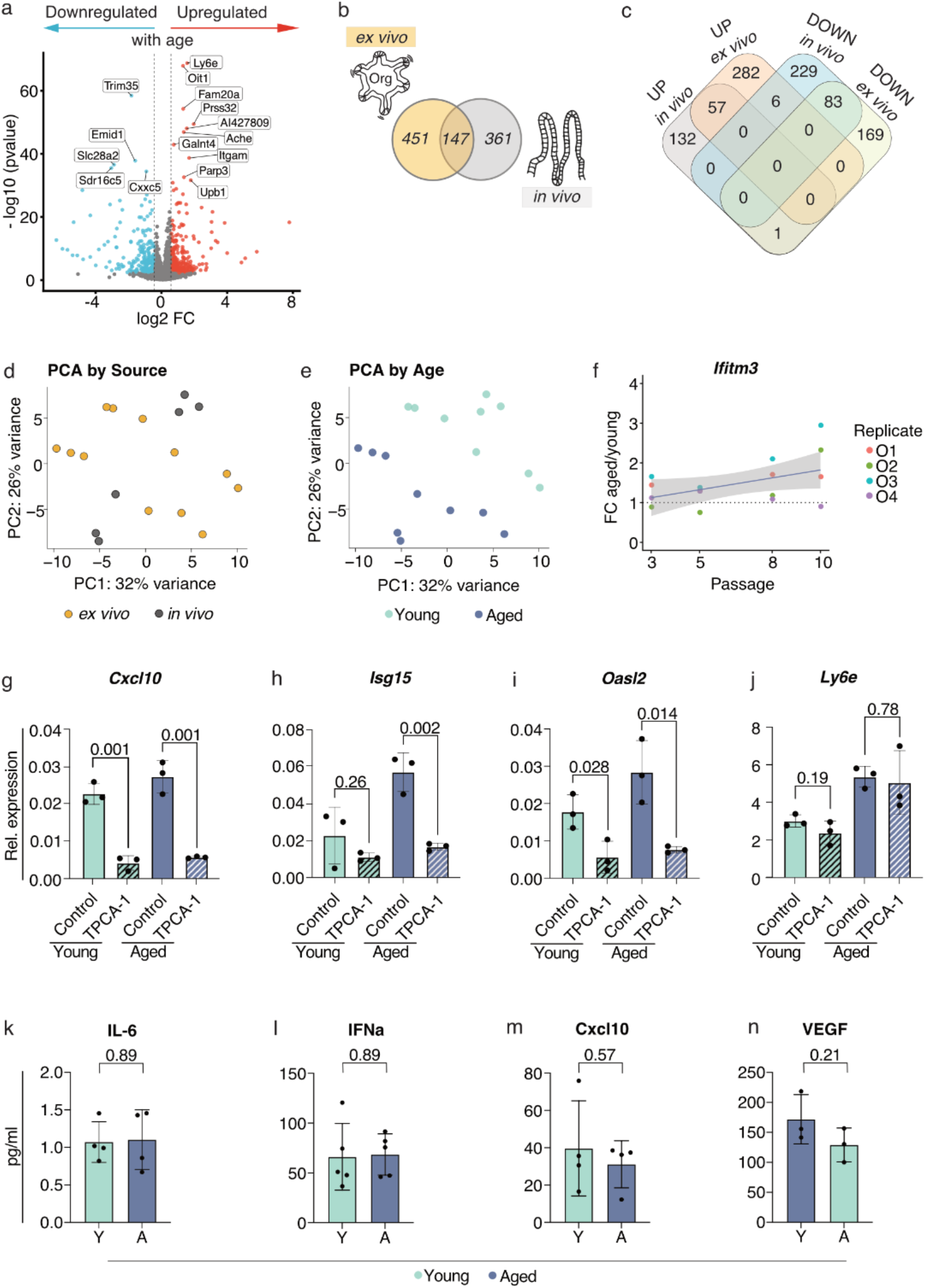
Comparison of intestinal organoids from young and aged mice by transcriptome and secretome. **a)** Volcano plot showing differentially expressed genes upon ageing (aged over young) in small intestinal organoids, red: significantly (FDR≤10%) upregulated (log2 FC > 0.5), blue: significantly (FDR≤10%) downregulated (log2 FC < (−0.5)), grey: not significant and absolute log2 FC < 0.); gene names of the 15 most significantly changed genes upon ageing are plotted, b) VENN diagram showing the overlap of 147 significantly (FDR≤10%) DEGs (absolute log2 FC > 0.5) as determined by bulk RNA-seq, between *ex vivo* cultured intestinal organoids, and *in vivo* intestinal epithelium, c) VENN diagram ofb) showing the overlap of significantly (FDR≤10%) upregulated (57) and downregulated (83) genes upon ageing between *ex vivo* cultured intestinal organoids and *in vivo* intestinal epithelium, **d)** + **e)** Intestinal organoids display a stable age effect in a joint analysis of bulk RNA-Seq data of young and aged *ex vivo* cultured intestinal organoids and *in vivo* intestinal epithelium. The color indicates in **d)** the source of the samples, in **e)** the age of the samples, f) Fold change (aged over young) expression of *Ifitm3* in intestinal organoids is over the time course of ten passages (eleven weeks of culture) constantly increased. Transcript levels were assessed in young and aged small intestinal organoids by qPCR (normalized to *36B4*) (n=4 per age), **g) - j)** Transcriptional changes of **g)** *CxcllO,* **h)** *Isg15*, **i)** *Oas12* & **j)** *Ly6e* upon 24 hour treatment with 5 μM TPCA-1, a NFkB & STAT3 signaling inhibitor, in young and aged intestinal organoids. Transcript levels were assessed in young and aged small intestinal organoids by qRT-PCR (normalized to *36B4*) (n=3 per age and per treatment), bars represent the average measure, data points represent the result of each biological replicate; statistical significance was tested by an unpaired t-test (two-sided) and p-Value is indicated on respective comparison; error bars indicate mean with standard deviation. **k-n)** Electrochemiluminescence-based cytokine assay to measure inflammation- and SASP-associated cytokines **k)** IL-6, **l)** IFNα, **m)** Cxcl10/IP-10, n) VEGF indicated no significant change in cytokine secretion between young and aged organoids. Medium of young and aged organoids was measured 2 days after medium renewal. Bars represent the average measure for all young (n=4) and all aged organoids (n=4), data points represent the result of each biological replicate, statistical significance was tested by an unpaired t-test (two-sided) and p-Value is indicated on respective comparison and error bars indicate mean with standard deviation; cytokine concentration is indicated in pg/ml (DEGs: differentially expressed genes, log2 FC: log_2_ fold change, SASP: senescence-associated secretory phenotype).

**Extended Data Figure 7:**
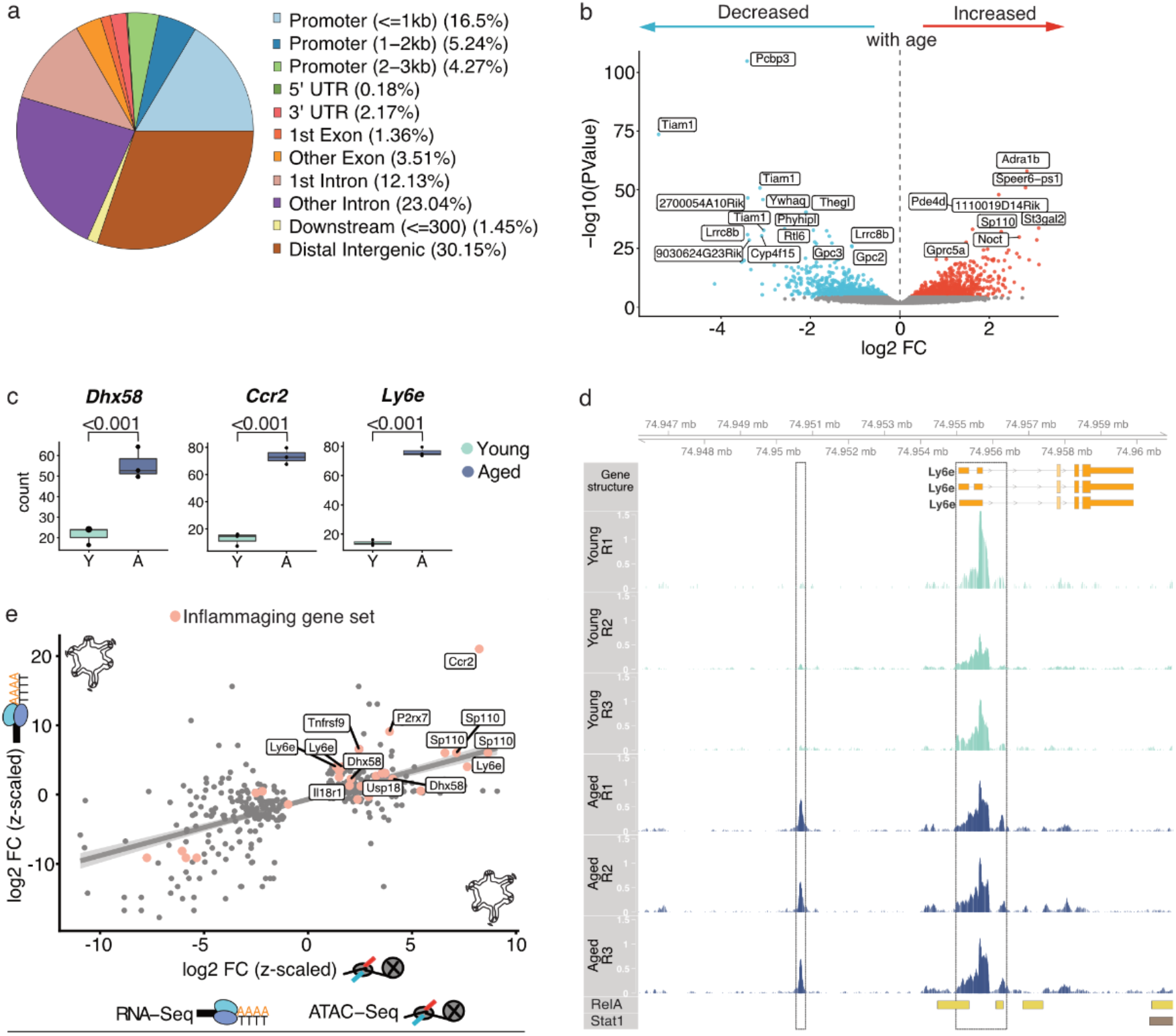
Changes in chromatin states accompany epithelial inflammaging. **a)** Pie chart represents the distribution of mapped regions of the ATAC-Seq experiment across genomic regions, **b)** Chromatin accessibility in intestinal epithelial cells undergoes changes upon ageing and persists in *ex vivo* culture. Volcano plot showing significantly (FDR≤10%) changed peak sizes indicative for chromatin accessibility of genetic loci upon ageing (aged over young) in small intestinal organoids derived from young (n=3) and aged (n=3) mice; 25 most significantly changed chromatin peaks are labeled, red: increased (log2 FC > 0), blue: reduced (log2 FC < 0), grey: not significantly changed, **c)** Chromatin accessibility for regulatory regions mapped to *Dhx58, Ccr2* and *Ly6e* is increased in intestinal organoids upon ageing. Count plots for *Dhx58, Ccr2*, and *Ly6e* for differentially open chromatin regions determined by ATAC-Seq. Bars represent the average counts for all young and all aged samples, data points represent the counts of the biological replicate, adjusted p-Value is indicated on respective comparison, **d)** Regulatory region assigned to *Ly6e* displays increased open chromatin upon ageing in the small intestinal epithelium. Peak density is shown for all young (n=3) and aged (n=3) samples in genomic coordinates close to *Ciita* by R/Gviz. RelA and Stat1 binding sites determined by LOLA are indicated at the bottom, **e)** Scatterplot of the effects of ageing on the expression of genes as determined by bulk RNA-seq in intestinal organoids (y-axis, z-scaled log2 FC) and on the chromatin accessibility as determined by ATAC-Seq in intestinal organoids (x-axis, z-scaled log2 FC). Genes of the inflammaging gene set are color-highlighted and a linear regression line indicates the agreement between the compared data sets (95 % confidence interval shaded grey). Only genes significantly (FDR≤10%) differently expressed in intestinal organoids are included (log2 FC: log_2_ fold change).

**Extended Data Figure 8:**
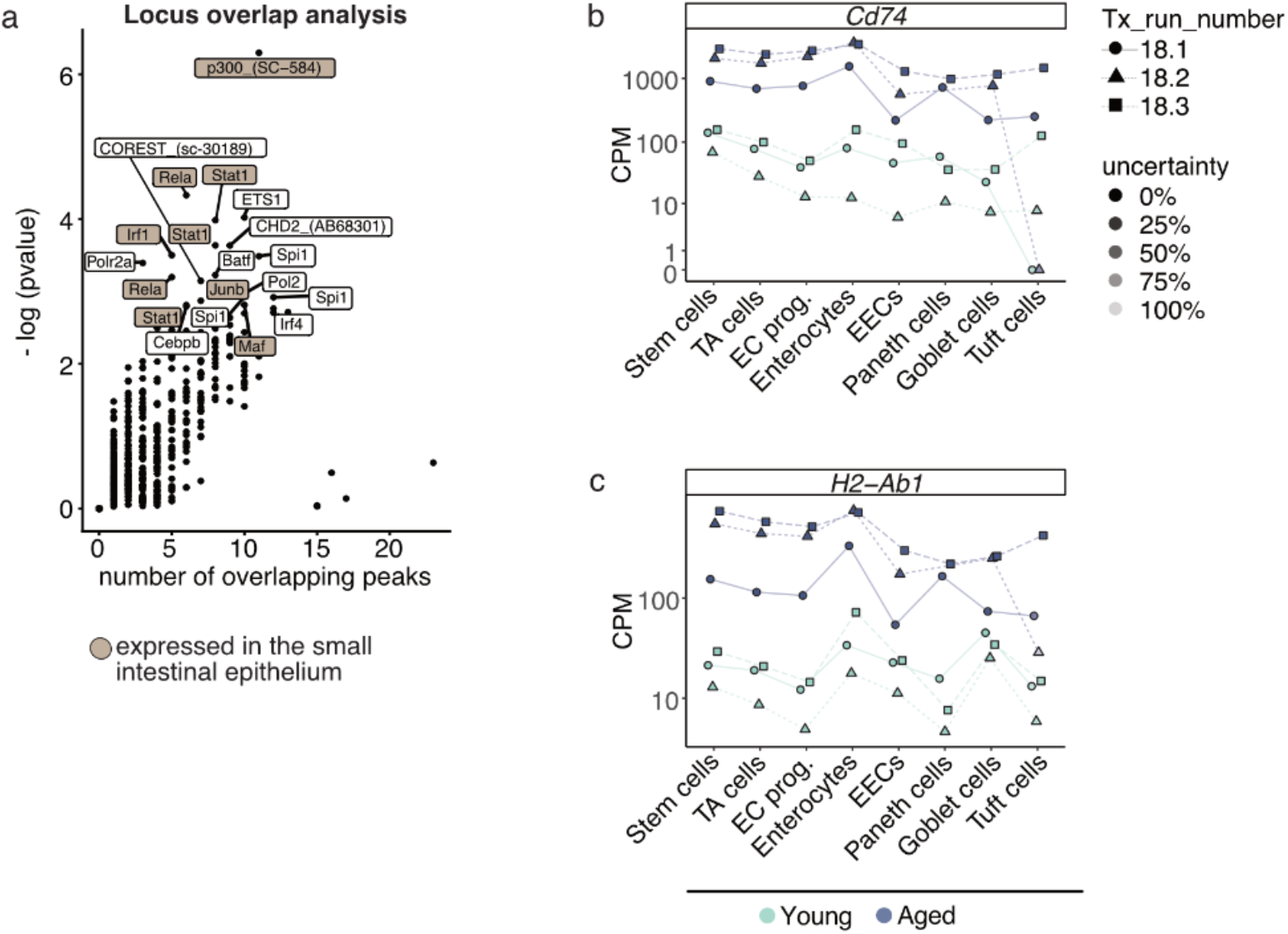
Transcription factors involved in age-related chromatin remodeling and MHC class II expression upon ageing. **a)** Locus Overlap Analysis (LOLA) determines enriched overlap between annotated transcription factor (TF) bindings sites and open chromatin regions of the ATAC-Seq experiment in intestinal organoids. Color highlighted are TFs that are expressed in the intestinal epithelium. Y-axis indicates the significance of overlap, the x-axis indicates overlap frequency, 20 most significant TF overlaps are labeled, **b) + c)** Expression of **MHC** class **II** encoding genes **b)** *Cd74* and **c)** *H2-Ab1* are upregulated in all cell types of the intestinal epithelium upon ageing, most robustly in Stem cells, TA cells, enterocyte progenitors, and enterocytes. CPM are plotted per cell type and per sample. The shape of the data points indicates the respective day of the experiment (10x library generation), transparency indicates the uncertainty of the respective data points and is given as the size of the 95% confidence interval (for the mean estimate across cells) relative to the plot height, color indicates age. (CPM: counts per million mapped reads, TA cells: transit-amplifying cells)

## Literature

1. SanMiguel, J. M., Young, K. & Trowbridge, J. J. Hand in hand: intrinsic and extrinsic drivers of aging and clonal hematopoiesis. Exp. Hematol. 91, 1–9 (2020).

2. Tyrrell, D. J. & Goldstein, D. R. Ageing and atherosclerosis: vascular intrinsic and extrinsic factors and potential role of IL-6. Nat. Rev. Cardiol. 18, 58–68 (2021).

3. Funk, M. C., Zhou, J. & Boutros, M. Ageing, metabolism and the intestine. EMBO Rep. 21, e50047 (2020).

4. Zheng, D., Liwinski, T. & Elinav, E. Interaction between microbiota and immunity in health and disease. Cell Res. 30, 492–506 (2020).

5. Pentinmikko, N. & Katajisto, P. The role of stem cell niche in intestinal aging. Mech. Ageing Dev. 191, 111330 (2020).

6. Barker, N. et al. Identification of stem cells in small intestine and colon by marker gene Lgr5. Nature 449, 1003–1007 (2007).

7. Beumer, J. & Clevers, H. Cell fate specification and differentiation in the adult mammalian intestine. Nat. Rev. Mol. Cell Biol. 22, 39–53 (2021).

8. Ayabe, T. et al. Secretion of microbicidal alpha-defensins by intestinal Paneth cells in response to bacteria. Nat. Immunol. 1, 113–118 (2000).

9. Biton, M. et al. T Helper Cell Cytokines Modulate Intestinal Stem Cell Renewal and Differentiation. Cell 175, 1307–1320.e22 (2018).

10. Guo, H., Gibson, S. A. & Ting, J. P. Y. Gut microbiota, NLR proteins, and intestinal homeostasis. J. Exp. Med. 217, (2020).

11. Bajic, D. et al. Gut Microbiota-Derived Propionate Regulates the Expression of Reg3 Mucosal Lectins and Ameliorates Experimental Colitis in Mice. J. Crohns. Colitis 14, 1462–1472 (2020).

12. Nalapareddy, K. et al. Canonical Wnt Signaling Ameliorates Aging of Intestinal Stem Cells. Cell Rep. 18, 2608–2621 (2017).

13. Mihaylova, M. M. et al. Fasting Activates Fatty Acid Oxidation to Enhance Intestinal Stem Cell Function during Homeostasis and Aging. Cell Stem Cell 22, 769–778.e4 (2018).

14. Pentinmikko, N. et al. Notum produced by Paneth cells attenuates regeneration of aged intestinal epithelium. Nature 571, 398–402 (2019).

15. Elderman, M. et al. The effect of age on the intestinal mucus thickness, microbiota composition and immunity in relation to sex in mice. PLoS One 12, e0184274 (2017).

16. Sovran, B. et al. Age-associated Impairment of the Mucus Barrier Function is Associated with Profound Changes in Microbiota and Immunity. Sci. Rep. 9, 1437 (2019).

17. Gebert, N. et al. Region-Specific Proteome Changes of the Intestinal Epithelium during Aging and Dietary Restriction. Cell Reports vol. 31 107565 (2020).

18. Furman, D. et al. Chronic inflammation in the etiology of disease across the life span. Nat. Med. 25, 1822–1832 (2019).

19. Kim, B.-H. et al. Interferon-induced guanylate-binding proteins in inflammasome activation and host defense. Nat. Immunol. 17, 481–489 (2016).

20. Haber, A. L. et al. A single-cell survey of the small intestinal epithelium. Nature 551, 333–339 (2017).

21. Sato, T. et al. Single Lgr5 stem cells build crypt-villus structures in vitro without a mesenchymal niche. Nature 459, 262–265 (2009).

22. Østvik, A. E. et al. Intestinal Epithelial Cells Express Immunomodulatory ISG15 During Active Ulcerative Colitis and Crohn’s Disease. J. Crohns. Colitis 14, 920–934 (2020).

23. Chi, X. et al. RORα is critical for mTORC1 activity in T cell-mediated colitis. Cell Reports vol. 36 109682 (2021).

24. Beaurivage, C. et al. Development of a human primary gut-on-a-chip to model inflammatory processes. Sci. Rep. 10, 21475 (2020).

25. Nan, J. et al. TPCA-1 is a direct dual inhibitor of STAT3 and NF-κB and regresses mutant EGFR-associated human non-small cell lung cancers. Mol. Cancer Ther. 13, 617–629 (2014).

26. Guerrero, A. et al. Cardiac glycosides are broad-spectrum senolytics. Nat Metab 1, 1074–1088 (2019).

27. Dabitao, D., Margolick, J. B., Lopez, J. & Bream, J. H. Multiplex measurement of proinflammatory cytokines in human serum: comparison of the Meso Scale Discovery electrochemiluminescence assay and the Cytometric Bead Array. J. Immunol. Methods 372, 71–77 (2011).

28. Benayoun, B. A. et al. Remodeling of epigenome and transcriptome landscapes with aging in mice reveals widespread induction of inflammatory responses. Genome Res. 29, 697–709 (2019).

29. Buenrostro, J. D., Wu, B., Chang, H. Y. & Greenleaf, W. J. ATAC-seq: A Method for Assaying Chromatin Accessibility Genome-Wide. Curr. Protoc. Mol. Biol. 109, 21.29.1–21.29.9 (2015).

30. Sheffield, N. C. & Bock, C. LOLA: enrichment analysis for genomic region sets and regulatory elements in R and Bioconductor. Bioinformatics 32, 587–589 (2016).

31. Neurath, M. F. Targeting immune cell circuits and trafficking in inflammatory bowel disease. Nat. Immunol. 20, 970–979 (2019).

32. Roche, P. A. & Furuta, K. The ins and outs of MHC class II-mediated antigen processing and presentation. Nat. Rev. Immunol. 15, 203–216 (2015).

33. Waldman, A. D., Fritz, J. M. & Lenardo, M. J. A guide to cancer immunotherapy: from T cell basic science to clinical practice. Nature Reviews Immunology vol. 20 651–668 (2020).

34. Smigiel, K. S., Srivastava, S., Stolley, J. M. & Campbell, D. J. Regulatory T-cell homeostasis: steady-state maintenance and modulation during inflammation. Immunol. Rev. 259, 40–59 (2014).

35. Lasry, A., Zinger, A. & Ben-Neriah, Y. Inflammatory networks underlying colorectal cancer. Nat. Immunol. 17, 230–240 (2016).

36. Kasler, H. & Verdin, E. How inflammaging diminishes adaptive immunity. Nature Aging vol. 1 24–25 (2021).

37. He, D. et al. Gut stem cell aging is driven by mTORC1 via a p38 MAPK-p53 pathway. Nature Communications vol. 11 (2020).

38. Moorefield, E. C. et al. Aging effects on intestinal homeostasis associated with expansion and dysfunction of intestinal epithelial stem cells. Aging 9, 1898–1915 (2017).

39. Tabula Muris Consortium. A single-cell transcriptomic atlas characterizes ageing tissues in the mouse. Nature 583, 590–595 (2020).

40. Krausgruber, T. et al. Structural cells are key regulators of organ-specific immune responses. Nature 583, 296–302 (2020).

41. Lu, J. et al. Characterization of an in vitro 3D intestinal organoid model by using massive RNAseq-based transcriptome profiling. Scientific Reports vol. 11 (2021).

42. Dobin, A. et al. STAR: ultrafast universal RNA-seq aligner. Bioinformatics 29, 15–21 (2013).

43. Ewels, P. A. et al. The nf-core framework for community-curated bioinformatics pipelines. Nat. Biotechnol. 38, 276–278 (2020).

44. Love, M. I., Huber, W. & Anders, S. Moderated estimation of fold change and dispersion for RNA-seq data with DESeq2. Genome Biol. 15, 550 (2014).

45. Korotkevich, G. et al. Fast gene set enrichment analysis. doi: 10.1101/060012.

46. Liberzon, A. et al. The Molecular Signatures Database (MSigDB) hallmark gene set collection. Cell Syst 1, 417–425 (2015).

47. Zheng, G. X. Y. et al. Massively parallel digital transcriptional profiling of single cells. Nat. Commun. 8, 14049 (2017).

48. Stuart, T. et al. Comprehensive Integration of Single-Cell Data. Cell vol. 177 1888–1902.e21 (2019).

49. Wickham, H. ggplot2: Elegant Graphics for Data Analysis. (Springer Science & Business Media, 2009).

50. Subramanian, A. et al. Gene set enrichment analysis: a knowledge-based approach for interpreting genome-wide expression profiles. Proc. Natl. Acad. Sci. U. S. A. 102, 15545–15550 (2005).

51. Tirosh, I. et al. Dissecting the multicellular ecosystem of metastatic melanoma by single-cell RNA-seq. Science 352, 189–196 (2016).

52. Hao, Y. et al. Integrated analysis of multimodal single-cell data. Cell 184, 3573–3587.e29 (2021).

53. Gontarz, P. et al. Comparison of differential accessibility analysis strategies for ATAC-seq data. Sci. Rep. 10, 10150 (2020).

54. Robinson, M. D., McCarthy, D. J. & Smyth, G. K. edgeR: a Bioconductor package for differential expression analysis of digital gene expression data. Bioinformatics 26, 139–140 (2010).

55. Yu, G., Wang, L.-G. & He, Q.-Y. ChIPseeker: an R/Bioconductor package for ChIP peak annotation, comparison and visualization. Bioinformatics 31, 2382–2383 (2015).

56. Hahne, F. & Ivanek, R. Visualizing Genomic Data Using Gviz and Bioconductor. Methods Mol. Biol. 1418, 335–351 (2016).

57. Lawrence, M. et al. Software for computing and annotating genomic ranges. PLoS Comput. Biol. 9, e1003118 (2013).

58. Bankhead, P. et al. QuPath: Open source software for digital pathology image analysis. Sci. Rep. 7, 16878 (2017).

59. Schindelin, J. et al. Fiji: an open-source platform for biological-image analysis. Nat. Methods 9, 676–682 (2012).

